# Rigosertib promotes anti-tumor immunity via autophagic degradation of PD-L1 in colorectal cancer cells

**DOI:** 10.1101/2023.02.07.527284

**Authors:** Xinyi Zhou, Dongliang Fu, Hang Yang, Chenqin Le, Yier Lu, Jingsun Wei, Yang Tang, Jiawei Zhang, Ying Yuan, Kefeng Ding, Qian Xiao

**Author notes:** Corresponding authors: Qian Xiao Kefeng Ding Ying Yuan. These authors contributed equally to this work (Xinyi Zhou, Dongliang Fu). Abbreviations: AMPK: AMP activated protein kinase; ATG: autophagy related gene; BafA1: bafilomycin A1; CHX: cycloheximide; CQ: chloroquine; CRC: colorectal cancer; CTL: cytotoxic T lymphocyte; CTLA-4: cytotoxic T-lymphocyte-associated protein 4; DC: dendritic cell; GFP: green fluorescent protein; HGFR: hepatocyte growth factor receptor; IFN- : interferon; LC3: microtubule associated protein 1γ light chain 3; mRFP: monomeric red fluorescent protein; mTOR: mammalian target of rapamycin; PARP: poly (ADP-ribose) polymerase; PD-1: programmed cell death protein 1; PD-L1: programmed cell death ligand 1; RAS: rat sarcoma viral oncogene homolog; RGS: rigosertib; ULK1: UNC-51-like kinase 1.

## Abstract

Rigosertib (RGS) is a benzyl styryl sulfone which exhibits impressive cytotoxicity in cancer cells. However, its modulating effect on tumor immune microenvironment remains elusive. In our experiments, compared with immunodeficient mouse model, increased tumor growth arrest and robust anti-tumor immunity were observed in RGS-treated colorectal cancer (CRC) xenograft tumors in immunocompetent mice.

Intriguingly, RGS markedly down-regulated programmed cell death ligand 1 (PD-L1) expression in both vivo and in vitro. Meanwhile, RGS increased autophagic vacuole number in CRC cells as seen by transmission electron microscopy and immunofluorescence. Moreover, increased LC3-II level and tandem- mRFP- GFP- LC3 labeled vacuole accumulation demonstrated RGS-induced autophagic flux.

Mechanistically, it is the activation of AMP-activated protein kinase-UNC-51-like kinase 1 (AMPK-ULK1) axis, rather than the canonical mTOR signaling pathway, that plays a pivotal role in RGS-induced autophagy. AMPK-ULK1 dependent autophagy inhibition, by either short interfering RNA or chemical inhibitors, blocked RGS-induced PD-L1 degradation. Finally, RGS exhibited synergistic anti-tumor activity with cytotoxic T-lymphocyte-associated protein 4 monoclonal antibody in the CRC xenograft model. Furthermore, apart from the immunomodulatory effect, we also confirmed the direct cytotoxicity of RGS in inducing mitochondria-related apoptosis. Altogether, considering its PD-L1 inhibitory and cytotoxic effects, RGS could be a promising drug for CRC therapy.

## Introduction

Colorectal cancer (CRC) is the fourth most common cause of cancer-related mortality in China [1]. The prognosis for metastatic CRC is poor, but the introduction of anti-programmed cell death protein 1 (PD-1) and anti-programmed cell death ligand 1 (PD-L1) monoclonal antibodies (mAbs) to its treatment has significantly improved survival of patients, especially patients with deficient DNA mismatch repair or microsatellite instability-high [2].

In the tumor microenvironment, cancer cells evade immune destruction by interacting with the host immune system via immune checkpoints, eventually allowing tumor cells to escape immune surveillance. PD-L1 is a transmembrane glycosylated protein that is often overexpressed in various cancers. As an immune checkpoint, it interacts with PD-1, a T cell inhibitory immune checkpoint receptor, to shut down the activity of cytotoxic T lymphocytes (CTLs) and help tumors escape eradication [3]. The regulation of the expression of PD-L1 is complex and occurs at different levels. For instance, interferon-γ (IFN-γ), which is secreted by activated T cells in the tumor microenvironment, induces the surface expression of PD-L1 on tumors *via* the Janus tyrosine kinase-signal transducer and activator of transcription pathway at the transcriptional level [4]. In addition, activation of intrinsic oncogenic pathways, such as rat sarcoma viral oncogene homolog (RAS) downstream RAF-MEK-ERK activation, contributes to the constitutive expression of PD-L1. Previously, Coelho et al. reported that the oncogenic RAS-RAF-MEK-ERK pathway increased the expression of PD-L1 by stabilizing PD-L1 mRNA, whereas inhibition of this oncogenic pathway via short interfering RNAs (siRNAs) or inhibitors reduced the expression of PD-L1 and enhanced anti-tumor immunity [5].

Autophagy is a highly conserved process, wherein eukaryotic cells deliver large aggregates of proteins and other cellular components to lysosomes for degradation. Based on the mechanisms involved, autophagy is categorized into macroautophagy, microautophagy, and chaperone-mediated autophagy [6]. Of these, macroautophagy/autophagy is the most well-studied process. Autophagy not only serves as an adaptive process induced by starvation, hypoxia, or innutrition, but also participates in degradation of pathologic proteins induced by pharmacological intervention [7]. Thus, numerous studies have been conducted with the aim of developing pharmacological modulators of autophagy to treat cancers.

Rigosertib (RGS), a benzyl styryl sulfone, acts as a non-ATP competitive multiple kinase inhibitor, suppressing the proliferation of various tumor cells both in vivo and in vitro [8]. Although the direct target of RGS remains controversial, the anti-tumor effect of RGS has been confirmed in various malignancies. Dai et al. reported that RGS inhibited the growth of diffuse large B-cell lymphoma through cytoplasmic sequestration of sumoylated CMYB- tumor necrosis factor receptor associated factor 6 complex [9]. Furthermore, Chapman et al. found that RGS was selectively cytotoxic towards chronic lymphocytic leukemia cells via inhibition of phosphoinositide 3-kinase (PI3K)-AKT pathway and induction of oxidative stress [10]. Recently, we found that RGS, as a RAS signaling disruptor, disrupted the activation of RAS-RAF-MEK-ERK and induced mitotic arrest and mitochondria-related apoptosis in RAS-mutated CRC cell lines [11]. Recently, Yan et al. found that, in addition to its direct cytotoxicity, RGS enriched dendritic cells (DCs) and induced the infiltration by activated T cells in the tumor microenvironment by inducing the up-regulation of CD40 in melanoma cells [12]. However, the modulating effect of RGS on PD-L1 expression remains unclear.

In our study, we demonstrated for the first time that RGS inhibited the expression of PD-L1 in CRC cells both in vivo and in vitro, and strengthened the anti-tumor immunity in the CT26 murine colorectal xenograft tumors. More importantly, we found that the RGS-induced suppression of PD-L1 was attributed, at least partially, to autophagy-mediated specific degradation. Meanwhile, our study confirmed the direct cytotoxicity of RGS in inducing mitochondria-related apoptosis. In summary, these data provided strong support for the use of RGS as a promising drug for the therapy of CRC owing to its targeting PD-L1 and its dual cytotoxic effect.

## Results

### RGS converted the immune environment in CRC from “cold” to “hot”

To investigate the potential role of RGS in anti-tumor immunity, we compared the anti-tumor effect of RGS in CT26 murine colorectal xenograft tumors based on immune competent (BALB/c) to that in immune deficient (nude) mice (Fig. S1A). Interestingly, after 10 days of treatment with RGS, significant tumor regression was only observed in BALB/c mice but not in those with immune deficiency (Fig. 1A, 1B and S1C). Other than the direct cytotoxicity of RGS in human CRC cells that we reported previously [11], the little arrest of tumor growth in nude mice revealed the resistance of murine CT26 cells to this cytotoxicity, and this resistance was also verified by the CCK8 assay in vitro (Fig. S1D). Accordingly, after treatment with RGS, no obvious change in apoptotic signal was observed in the xenograft tumors in nude mice; however, in BALB/c mice, stronger fluorescence of clustered cleaved caspase 3 was observed in the RGS-treated CT26 tumors compared to phosphate buffered saline (PBS)-treated mice (Fig. 1C and data not shown). Thus, the differences in tumor growth curves and apoptotic signal change between immune competent and immune deficient xenograft mouse models suggested that, besides the direct cytotoxicity of RGS, the anti-tumor effect of RGS was closely related to immune response.

**Figure 1.**
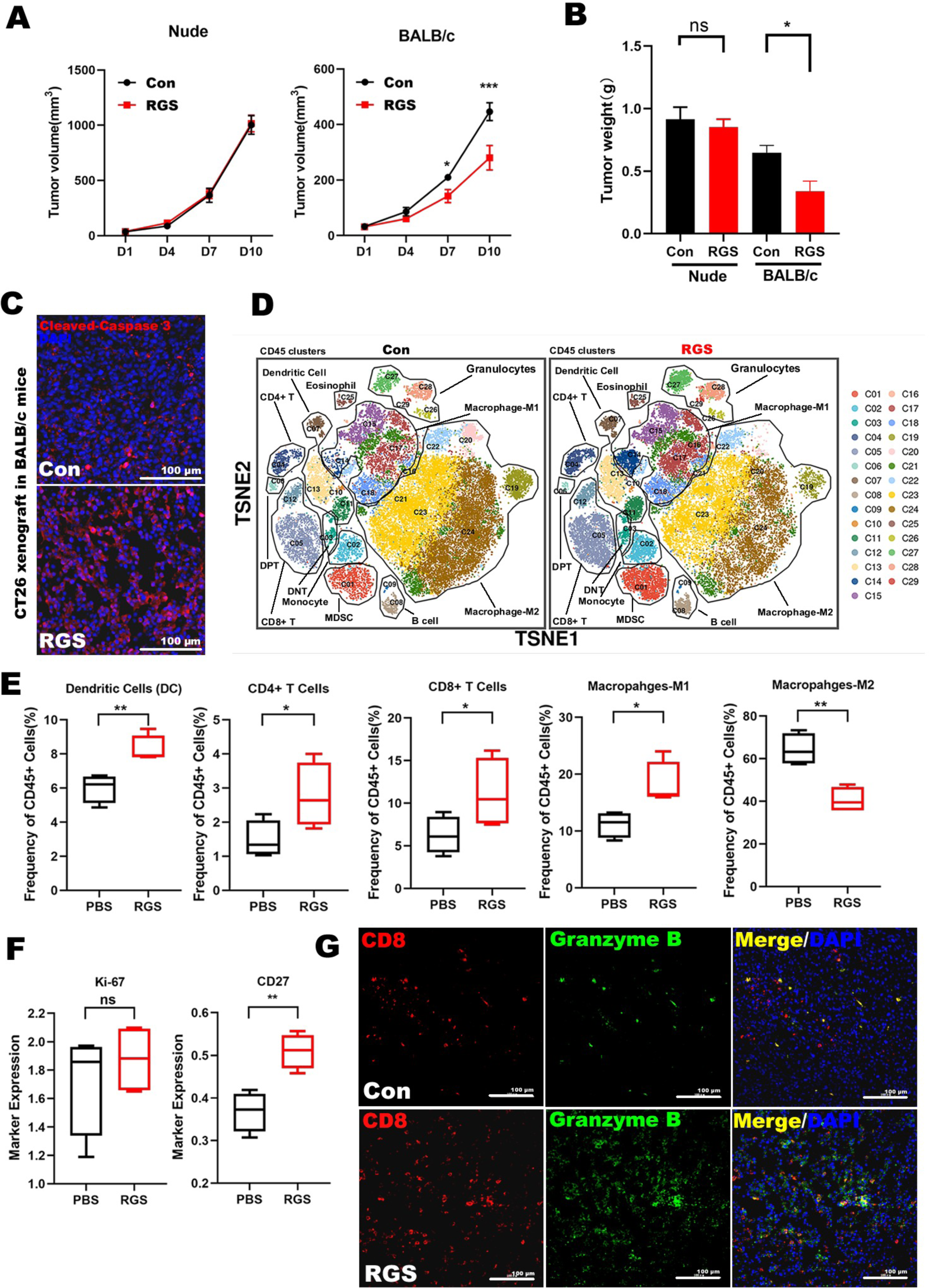
RGS converted the immune environment of colorectal cancer from “cold” to “hot.” (**A and B**) Anti-tumor effect of RGS was assessed in CT26 xenograft models based on nude or BALB/c mice separately. Tumor volumes were measured every 3 d after palpable tumors reached 50–100 mm^3^. Tumor growth and tumor weight were significantly suppressed in the RGS-treated CT26 xenograft tumors based on immune-competent BALB/c mice (N = 3–5). (**C**) Immunostaining of cleaved caspase 3 in the CT26 xenograft tumors based on immune competent BALB/c mice. Scale bar, 100 μm. (**D**) t-SNE visualization of CD45+ tumor-infiltrating leukocytes of CT26 xenograft tumors from BALB/c mice showed a total of 29 clusters and the distributions in clusters of the PBS or RGS-treated tumors. (**E**) Data derived from CyTOF. Frequencies of DCs, CD4+ T cells, CD8+ T cells, and macrophages between PBS and RGS-treated tumors. (**F**) Differences in expression of Ki-67 and CD27 between PBS and RGS-treated tumors. Data shown are mean ± SEM. \**P* < 0.05; \*\**P* < 0.01; \*\*\**P* < 0.001. (**G**) Immunostaining of CD8 (CTLs marker) and granzyme B (activity of CTLs) in the CT26 xenograft tumors based on BALB/c mice. Scale bar, 100 μm.

To explore how RGS affected the tumor immune environment, we investigated the immune landscape of CD45+ tumor-infiltrating leukocytes in CT26 xenograft tumors from BALB/c mice by mass cytometry (CyTOF). A total of 41 biomarkers, including cell membrane proteins and functional molecules, were enrolled, and 29 clusters were identified (Fig. 1D and S1E). We identified the following major cell subsets: DCs, CD8+ T cells, CD4+ T cells, B cells, macrophages, monocytes, myeloid-derived suppressor cells along with a small population of granulocytes based on the expression levels of lineage markers (Fig. 1D, 1E, S1E and S1F). Compared to the control group, RGS induced significant expansion of T cells and B cells while no obvious natural killer cells cluster was observed in both groups. We found that both the populations of CD4+ T helpers (C04) and CD8+ CTLs (C05 and C12), particularly the CD69-positive activated CD8+ T cells (C05), were elevated in the RGS-treated tumors. While the CD4+ and CD8+ tumor-infiltrating T cells increased, we observed no remarkable alteration in the expression of CD25, an immune regulatory molecule mainly expressed on CD4+ regulatory T cells (Fig. S2A). Moreover, we found a prominent increase in the number of total tumor-infiltrating DCs (C07, C10, C11 and C13), the most efficient antigen presenting cells, especially the CD69+ CD80+ plasmocytoid DCs (C11). Among the myeloid populations, the percentages of granulocytes did not change, and the frequency of monocytes (C02), M1 phenotype macrophages (C14, C15, C17 and C18) and myeloid-derived suppressor cells (C01) increased following RGS treatment, while a 25% decrease of CD206+ M2 macrophages was observed in RGS-treated tumors (C19–C24). Meanwhile, RGS treatment showed little cytotoxicity towards the total tumor-infiltrating leukocytes, as measured by the expression level of Ki-67, a classical proliferation marker (Fig. 1F).

The expression of functional molecules in the leukocytes was also evaluated. Despite the little change in PD-1 expression (Fig. S2C), we found that the expression of CD27, another transmembrane co-stimulatory phosphor-glycoprotein mainly expressed on CD8+ cytotoxic T cells [13] (Fig. S2B), increased by 40% in the RGS-treated tumors (Fig. 1F). Meanwhile, as a membrane-associated ecto-enzyme involved in the immunosuppressive adenosinergic pathway [14], the expression of CD38 was evidently suppressed in M2 macrophages after RGS treatment (Fig. S2D). Considering that CD8+ CTLs, the major effector of anti-tumor immunity, could eliminate cancer cells by secreting granzyme B, an apoptotic trigger, we then examined the activity of CD8+ CTLs by measuring the levels of granzyme B. Subsequently, we found that treatment with RGS, indeed, increased the infiltration by CD8+ CTLs and release of granzyme B (Fig. 1G).

Together, these results suggested that RGS could reprogram the tumor immune microenvironment and increase activity of CTLs, which may contribute to its anti-tumor effect.

### RGS down-regulated PD-L1 in CRC cell lines both in vivo and in vitro

In the tumor immune microenvironment, activity of CTLs was negatively regulated by interactions between immune checkpoints and their ligands such as PD-1 and PD-L1 axis. Considering the increased activity of CTLs and robust expression of PD-1 in RGS-treated CT26 xenograft tumors (Fig. S2C), we aimed to determine whether RGS increased activity of CTLs by reducing levels of PD-L1 in tumor. Interestingly, lower levels of PD-L1 were, indeed, observed in RGS-treated CT26 tumors and cells both in vivo and in vitro (Fig. 2A and 2B).

**Figure 2.**
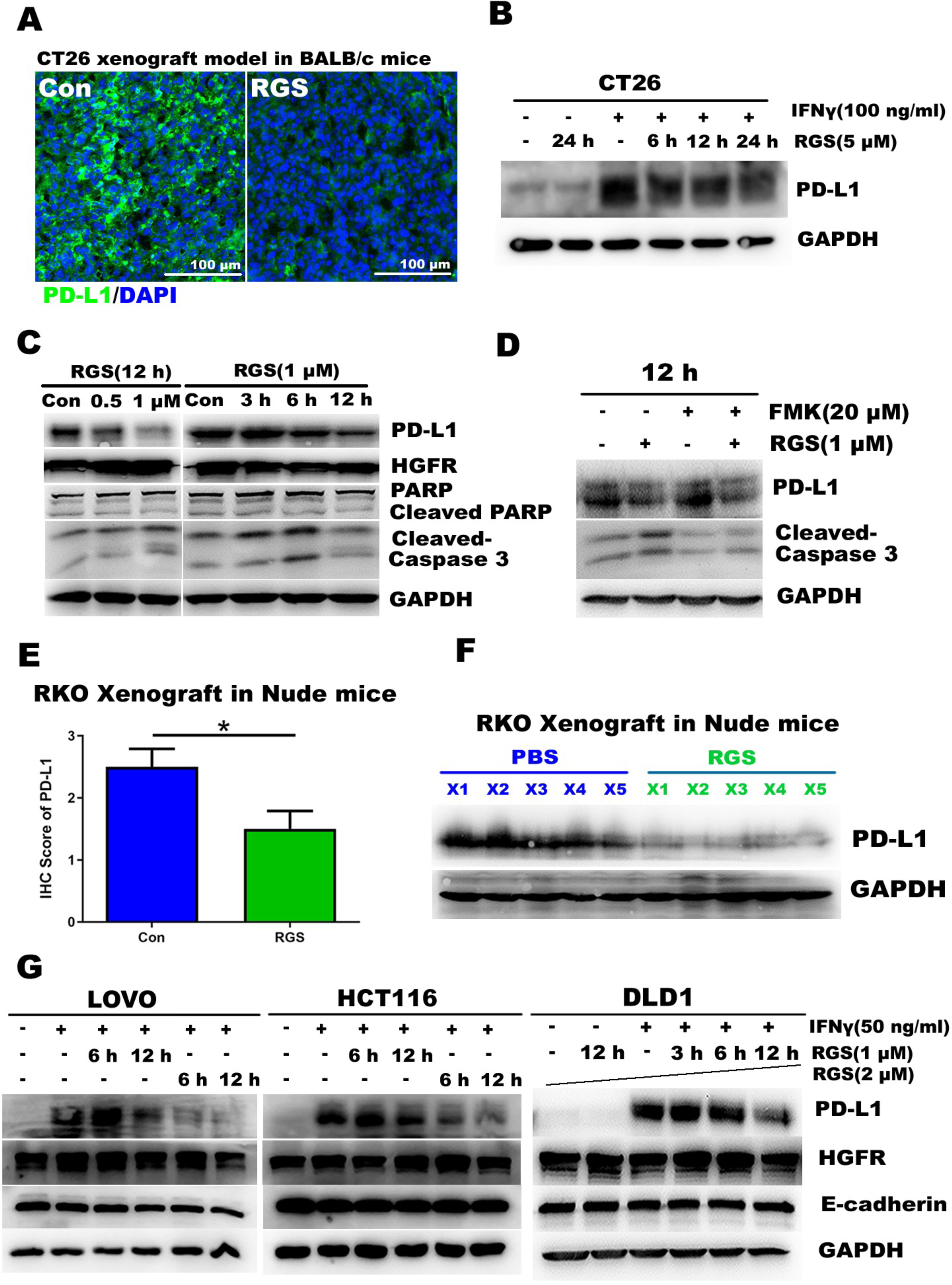
RGS down-regulated PD-L1 in colorectal cancer cell lines both in vivo and in vitro. (**A**) Immunostaining of PD-L1 in the CT26 xenograft tumors based on BALB/c mice. Scale bar, 100 μm. (**B**) Following incubation of CT26 cells with IFN-γ for 48 h, CT26 cells were treated with 5 μM RGS for various durations prior to lysis; the levels of inducible PD-L1 were examined by western blotting. (**C**) RKO cells M for various durations prior to μ lysis. PD-L1 and apoptosis related protein markers were examined as indicated. (**D**) RKO cells were treated with RGS, with or without pretreatment with Z-VAD-FMK, for 12 h prior to lysis. PD-L1 and apoptosis-related protein markers were measured by western blot analysis. (**E**) After intraperitoneal injection of RGS, the levels of PD-L1 in the subcutaneous RKO xenograft tumors were evaluated by immunohistochemical staining, and the immunoreactive scores were calculated. Data are presented as the mean ± SEM from three different experiments performed in triplicate. \**P* < 0.05. (**F**) The level of PD-L1 in subcutaneous RKO xenograft tumors were examined by western blotting. (**G**) After incubation with IFN-γ for 48 h, DLD1, LOVO, and HCT116 cells were treated with 1 or 2 μM RGS for various time periods prior to lysis. The levels of inducible PD-L1 and those of two other membrane proteins (HGFR and E-cadherin) were examined by western blotting.

To screen for a suitable human CRC cell line for investigating the mechanisms underlying the modulating effect of RGS on PD-L1 expression, we investigated the expression of PD-L1 protein in nine human CRC cell lines (HT29, DLD1, HCT116, SW480, SW620, RKO, LOVO, SW48, and Caco-2) by western blotting. The results showed that PD-L1 was highly expressed in RKO cells (Fig. S3A). Additionally, we found that, after RGS treatment, PD-L1 was down-regulated in RKO cells in a time- and dose- dependent manner (Fig. 2C). In contrast to murine CT26 cells, RKO cells were sensitive to the direct cytotoxicity of RGS. More specifically, we observed that incubation with RGS for 24 h markedly inhibited viability of RKO cells (Fig. S3B). Moreover, apoptosis-related proteins, such as cleaved caspase 3, cleaved caspase 9, and cleaved poly (ADP-ribose) polymerase (PARP), increased evidently after RGS treatment (Fig. S3C). By separating cytoplasmic and mitochondrial fractions, we found that the proapoptotic protein, Bcl-2-associated X protein, was significantly down-regulated in the cytoplasm after 24 h of exposure to RGS, and cytochrome c was significantly released from the mitochondria to the cytoplasm (Fig. S3D), indicating that RGS might induce mitochondria-related apoptosis. To further exclude the correlation between the PD-L1 inhibitory effect and the RGS-induced direct cytotoxicity, we determined the levels of cleaved caspase 3 and cleaved PARP in parallel experiments. We found that incubation with RGS for less than 12 h markedly decreased the expression of PD-L1; however, this short-term incubation did not notably increase the expression of apoptosis-related protein markers (Fig. 2C, similar to the results shown in Fig. S3C). Moreover, we noticed that pretreatment with the pan-caspase inhibitor, Z-VAD-FMK, did not rescue the RGS-induced decrease in the expression of PD-L1 (Fig. 2D). Therefore, we assumed that the RGS-induced reduction in the level of PD-L1 was not a result of its cytotoxicity.

Next, we aimed to confirm the PD-L1-reducing effect of RGS in RKO subcutaneous xenograft tumors. Similar to our previous findings [11], a significant inhibition of tumor weight and growth in the RGS-treated nude mice was observed, while no apparent body weight loss was observed (Fig. S3E–H). Furthermore, through immunohistochemistry and western blotting, we evaluated the level of PD-L1 in the subcutaneous RKO xenograft tumors and found that intraperitoneal administration of RGS significantly decreased the level of PD-L1 in vivo (Fig. 2E, 2F and S3I).

We further explored whether RGS could down-regulate the level of inducible PD-L1. As shown in Fig. 2G, similar to that in murine CT26 CRC cells, IFN-γ promoted the expression of PD-L1 in LOVO, HCT116, and DLD1 CRC cells, and PD-L1 levels were remarkably inhibited by RGS in a time- and dose-dependent manner. On the other hand, the levels of two other membrane proteins, hepatocyte growth factor receptor (HGFR) and E-cadherin, which are highly expressed in CRC cells, did not decrease, indicating that RGS down-regulated the expression of PD-L1 specifically.

Collectively, our results indicated that RGS decreased the expression of PD-L1 in CRC cells, in a process that was independent of its direct cytotoxic effect.

### The lysosome pathway participated in the RGS-induced inhibition of PD-L1

To explore the mechanism underlying the RGS-induced down-regulation of PD-L1 expression, we first examined whether the RGS-induced suppression of PD-L1 occurred at the transcriptional level. Real-time PCR analysis showed that the level of PD-L1 mRNA was not significantly altered after treatment with RGS (Fig. 3A).

**Figure 3.**
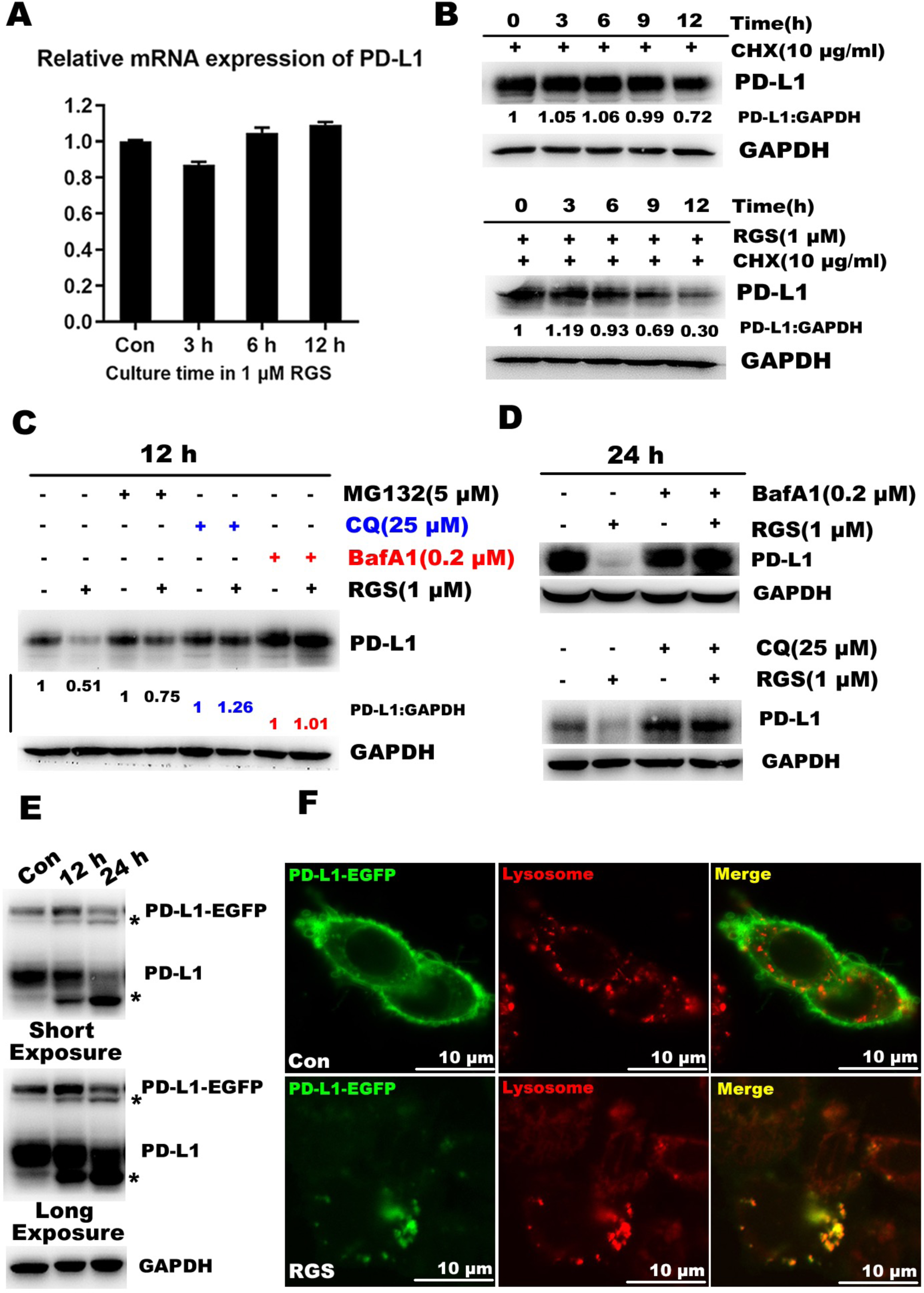
RGS accelerated the degradation of PD-L1 via the lysosomal pathway. (**A**) RKO cells were exposed to 1 μ RGS for 0–12 h. Total RNA was extracted and the expression of PD-L1 mRNA was analyzed by RT-PCR, using *Gapdh* as control. (**B**) RKO cells were treated with CHX in the presence or absence of RGS (1 μ different durations, and the levels of PD-L1 were then measured by western blotting. (**C and D**) RKO cells were pretreated with MG132, CQ, or BafA1 for 30 min to inhibit the function of proteasomes and lysosomes. Cells were then treated with or without RGS (1 μM) for 12–24 h, after which the levels of PD-L1 were determined. (**E**) In PD-L1-EGFP-RKO cells, the levels of PD-L1-EGFP and PD-L1 were determined after treatment with 1 μM RGS for 12–24 h; PD-L1 fragments are indicated by stars. (**F**) In PD-L1-EGFP-RKO cells, the lysosomes were stained using Lyso-tracker. After treatment with RGS (1 μ) for 12 h, the distribution of PD-L1-EGFP and lysosomes was determined under a confocal microscope. Scale bar, 10 μm.

Accordingly, we found that RGS treatment did not suppress the activation of RAF-MEK-ERK pathway, which plays a pivotal role in enhancing the transcription of PD-L1 (Fig. S4A). A recent study has shown that PD-L1 is secreted from cancer cells and suppresses the function of CD8+ T cells [15]. By detecting the level of secreted PD-L1 in culture media via ELISA, we found that RGS did not affect the secretion of PD-L1 (Fig. S4B). On the basis of these findings, we hypothesized that a degradation mechanism might be involved in the RGS-induced reduction of PD-L1 expression. To investigate this, we employed cycloheximide (CHX), a protein synthesis inhibitor, to block de novo protein synthesis; thus, any change in the level of PD-L1 would primarily reflect protein degradation. We exposed RKO cells to CHX in the presence or absence of RGS for different time points and measured the expression of PD-L1.

As shown in Fig. 3B, the results showed that the level of PD-L1 in RGS-treated cells decreased in a time-dependent manner. Meanwhile, the immunoblot bands corresponding to degraded PD-L1 fractions were elevated in a dose- and time-dependent manner after treatment with 0.5–1 μ RGS for 12–24 h (Fig. S4C).

Thus, these results indicated that RGS decreased the expression of PD-L1 by accelerating its degradation.

Lysosomal and proteasomal degradation are two of the major pathways involved in cellular protein turnover. We pretreated RKO cells with MG132 (a proteasome inhibitor), chloroquine (CQ), and bafilomycin A1 (BafA1) (two lysosome inhibitors) to block the proteasome and lysosome activities separately. Unlike MG132 pretreatment, CQ and BafA1 pretreatment completely abrogated the RGS-induced reduction in PD-L1 expression (Fig. 3C and 3D). To confirm that RGS-induced degradation of PD-L1 occurs in lysosomes, we constructed RKO cells that stably expressed the PD-L1-enhanced green fluorescent protein (EGFP) fusion protein. We found that, similar to the expression of endogenous protein, the expression of PD-L1-EGFP decreased in a time-dependent manner after RGS treatment (Fig. 3E).

Meanwhile, as shown in Fig. 3F, we detected that RGS treatment increased the distribution of PD-L1 to lysosomes.

Collectively, these results demonstrated the role of RGS in mediating the lysosomal degradation of PD-L1.

### RGS induced autophagy in CRC cells via AMP-activated protein kinase-UNC-51-like kinase 1 (AMPK-ULK1) axis

Previous studies have demonstrated that the major route for the delivery of proteins and organelles into lysosomes is through autophagy [16]. Therefore, we explored whether RGS could modulate autophagy in RKO cells. First, we determined the presence of autophagic vacuoles by transmission electron microscopy. As shown in Fig. 4A, we observed typical autophagic structures, including double-membrane structures containing cytoplasmic materials (autophagosomes) and single-membrane structures containing degraded cytoplasmic contents (autolysosomes), in RGS-treated RKO cells.

**Figure 4.**
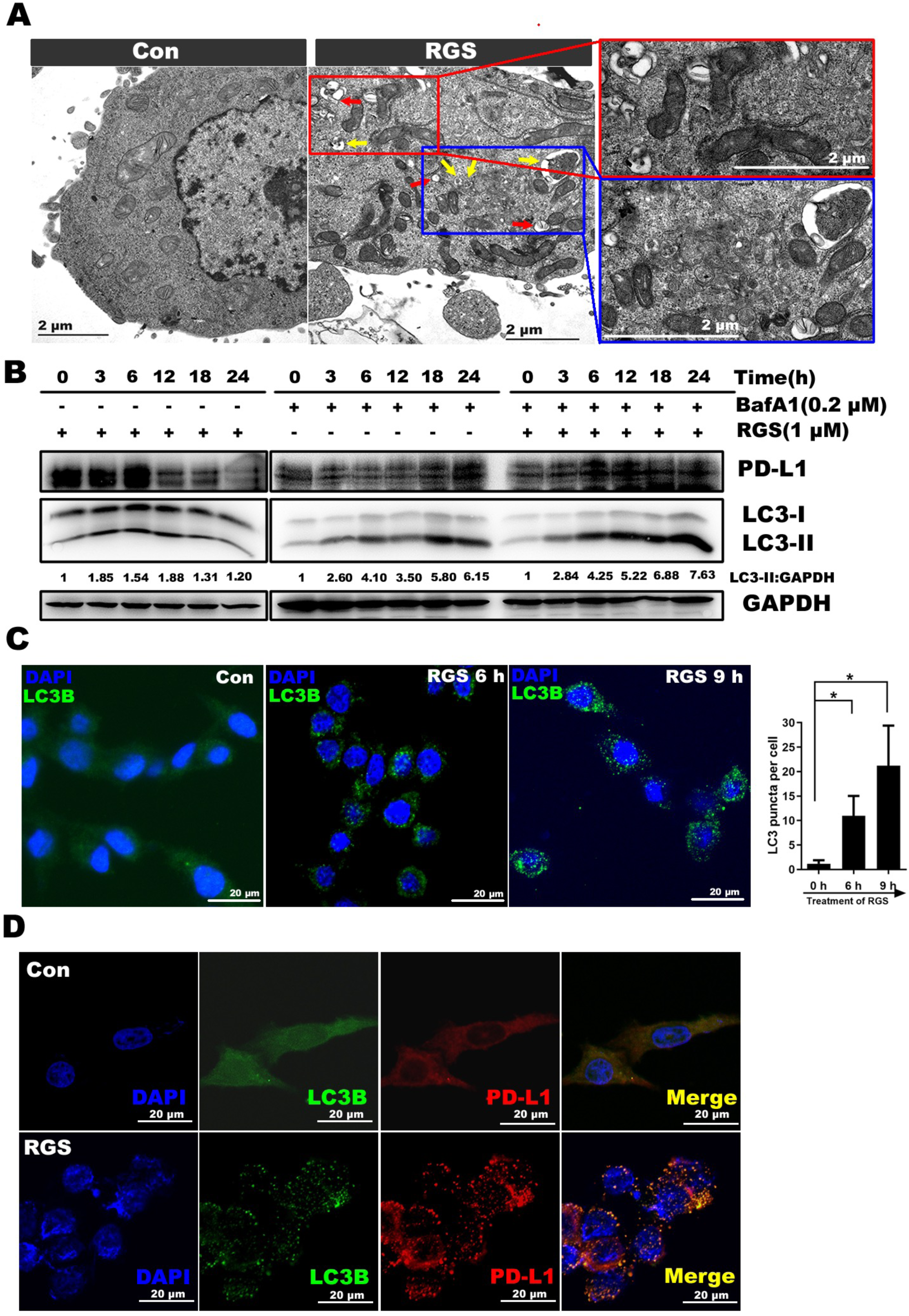
RGS induced autophagy in RKO cells. (**A**) Ultrastructural features of RKO cells with or without treatment with RGS (1 μ for 12 h were analyzed by electron microscopy. Typical images of autophagosomes (RED arrows) and autolysosomes (Yellow arrows) were shown at higher magnification. Scale bar, 2 μm. (**B**) RKO cells were pretreated with BafA1 for 30 min to inhibit lysosome function. Cells were then M) for 0–24 h, after which the levels of PD-L1 and μ LC3 were determined as indicated; levels of LC3-II and GAPDH were quantitated using Image J software. (**C**) RKO cells were exposed to RGS (1 μ) for 0–9 h, and then the distribution of LC3 was analyzed by immunofluorescence. Scale bar, 20 μm. For quantitative analysis, the number of LC3 puncta per cell was counted. At least two random fields from one section and two sections were analyzed in each independent experiment. The bar chart represents the mean ± SEM values from at least three independent experiments; at least 70 cells were counted in each group. \**P* < 0.05. (**D**) RKO cells were exposed to RGS (1 μ) for 9 h, and then the distribution of PD-L1 and LC3 was analyzed under a confocal microscope. Scale bar, 20 μm.

Next, we examined the turnover of microtubule associated protein 1 light chain 3 (LC3), a widely accepted autophagic marker, to evaluate the degree of autophagy in RGS-treated cells. During the formation of autophagosomes, LC3-I is converted to lipidated LC3-II, indicating the induction of autophagy. As shown in Fig. S5A and S5B, we noticed that RGS increased the expression of LC3-II in a time- and dose-dependent manner. Additionally, as the increased LC3-II is then degraded in autolysosomes during later stages of autophagy, pretreating cells with lysosome inhibitors to assess autophagic flux is recommended [7]. As shown in Fig. 4B, RGS increased the level of LC3-II until 12 h after incubation. When we prolonged incubation to 18–24 h, the level of LC3-II decreased, probably due to degradation of LC3-II in autolysosomes (Fig. 4B left). We thus employed BafA1 pretreatment to block any lysosome activity before RGS treatment. We found that BafA1 inhibited the RGS-induced degradation of PD-L1. Moreover, a larger accumulation of LC3-II was observed in the BafA1 + RGS group compared with that in the BafA1 group, indicating the induction of autophagic flux by RGS (Fig. 4B middle and right).

Concomitantly, pretreatment with CQ also indicated the induction of autophagic flux by RGS (Fig. S5C).

Furthermore, we assessed the number of autophagic vacuoles in RKO cells by LC3 immunofluorescence staining (Fig. 4C). Quantitative analysis showed increased numbers of LC3 puncta in RKO cells after 6–9 h of incubation with RGS. Moreover, as shown in Fig. 4D, these LC3-labeled vacuoles were confirmed to engulf PD-L1 in RGS-treated RKO cells. PD-L1 was mainly co-localized with LC3, whereas PD-L1 was localized in the cytoplasm and cellular membrane in untreated cells.

Furthermore, tandem fluorescent-tagged monomeric red fluorescent protein (mRFP)-green fluorescent protein (GFP)-LC3 was used to visualize the autophagic flux. Concretely, in autophagosomes, the tandem fluorescent-tagged LC3 shows both mRFP and GFP signals; whereas in autolysosomes, GFP fluorescence is relative sensitive to the acidic conditions, whereas mRFP is more stable [17]. We found that RGS evidently increased the fluorescent-tagged LC3 puncta, wherein the fluorescence intensity of mRFP was much stronger than GFP, indicating that the LC3-labeled autophagosomes fused with lysosomes after 12 h of incubation. Subsequently, blocking the lysosome activity using BafA1 restored the GFP signals and resulted in accumulation of LC3 puncta. Furthermore, compared with the BafA1-treated CRC cells, a larger accumulation of LC3 puncta was observed in the RGS + BafA1 group, which confirmed the induction of autophagic flux induced by RGS (Fig. S5D).

The activation of ULK1, a homologue of yeast autophagy related gene 1 (ATG1) serine/threonine protein kinase, promotes the initiation and maturation of autophagy [18]. The activity of ULK1 is tightly regulated by upstream AMPK and the mammalian target of rapamycin (mTOR); specifically, AMPK promotes autophagy by directly activating ULK1 through phosphorylation of Ser555 [19], while activated mTOR prevents ULK1 activation by phosphorylating ULK1 Ser757 [20]. As shown in Fig. 5A, no suppression in mTOR phosphorylation at Ser2448 occurred despite the slight changes in AKT phosphorylation, indicating that RGS could not inhibit mTOR signaling. Accordingly, we observed no obvious alteration in the phosphorylation of ULK1 at Ser757, while activated AMPK (phosphor-AMPK at Thr172) and ULK1 (Ser555) increased in a time- and dose-dependent manner (Fig. 5B–D). To clarify whether RGS-induced autophagy depended on AMPK-ULK1 activation, we used small-molecule inhibitors of AMPK-ULK1 axis. Blocking the upstream AMPK by Compound C restrained both the phosphorylation of AMPK and ULK1 induced by RGS; while SBI0206965, a ULK1 small molecule inhibitor [21], only suppressed the activation of downstream ULK1, not the AMPK kinase (Fig. 5E). Moreover, both Compound C and SBI0206965 blocked the increasing LC3-II level and LC3 fluorescence puncta induced by RGS (Fig. 5F and S5D).

**Figure 5.**
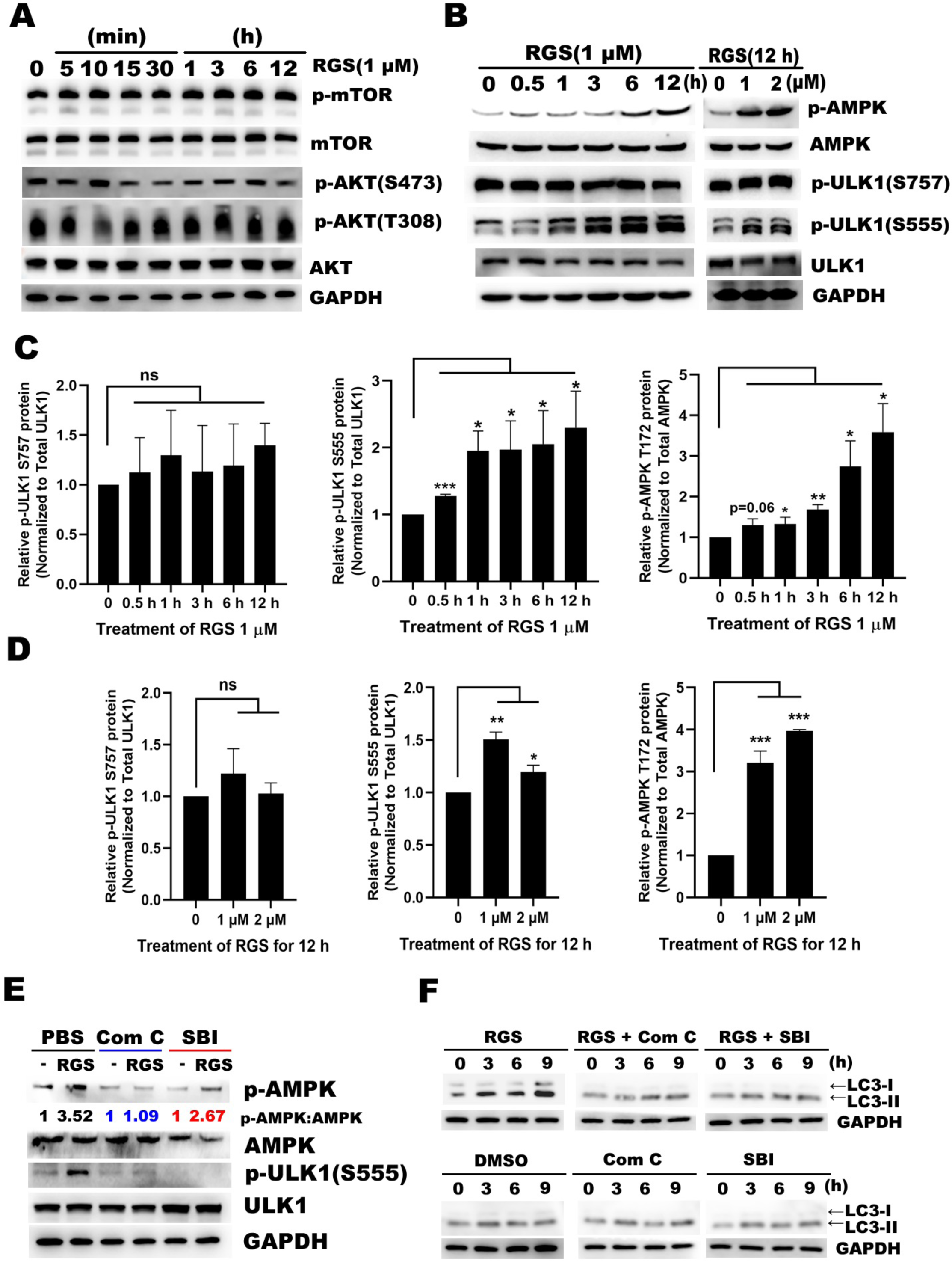
RGS induced autophagy in RKO cells by activating AMPK-ULK1 axis. (**A**) RKO cells were treated as indicated with 1 μ examined for levels of mTOR, p-mTOR (S2448), AKT, p-AKT (S473), and p-AKT (T308) by western blotting. (**B-D**) RKO cells were treated with 1 or 2 μ RGS for 12 M for various times prior to lysis. Lysates were examined for levels of AMPK, μ p-AMPK (T172), ULK1, p-ULK1 (S757), and p-ULK1 (S555) by western blotting. Data shown are mean ± SEM. \**P* < 0.05; \*\**P* < 0.01; \*\*\**P* < 0.001. (**E**) RKO cells were pretreated with Compound C and SBI0206965 for 30 min to inhibit the activity of AMPK and ULK1. Cells were then treated with or without RGS (1 μM) for 12 h, after which the levels of AMPK, p-AMPK (T172), ULK1, and p-ULK1 (S555) were determined. (**F**) RKO cells were pretreated with Compound C and SBI0206965 for 30 min. Cells were then treated with or without RGS (1 μ) for 0–9 h, after which the level of LC3 were determined as indicated.

These results suggested that the AMPK-ULK1 axis dependent autophagy was potentiated during treatment with RGS, which might participate in degrading PD-L1.

### Autophagy regulated the RGS-induced degradation of PD-L1

Besides blocking autophagy, inhibition of AMPK-ULK1 axis by pretreatment with Compound C, SBI0206965, and MRT68921 (another ULK1 inhibitor) [22] suppressed the degradation of PD-L1 induced by RGS, which indicated the dependence of PD-L1 degradation on AMPK-ULK1 dependent autophagy (Fig. 6A and 6B). In addition, we found that siRNA-mediated depletion of *ULK1* expression inhibited this degradative process (Fig. S6A). Two ubiquitin-like conjugation systems, the ATG12–ATG5 conjugate and LC3 systems, are tightly associated with the expansion of autophagosomes. As such, knocking down these two systems (using si*ATG5* and si*LC3*) indeed suppressed the degradation of PD-L1 induced by RGS (Fig. 6C and 6D). Furthermore, we found that rapamycin, a typical autophagic agonist, decreased the levels of PD-L1 in a time- and dose-dependent manner (Fig. S6B).

**Figure 6.**
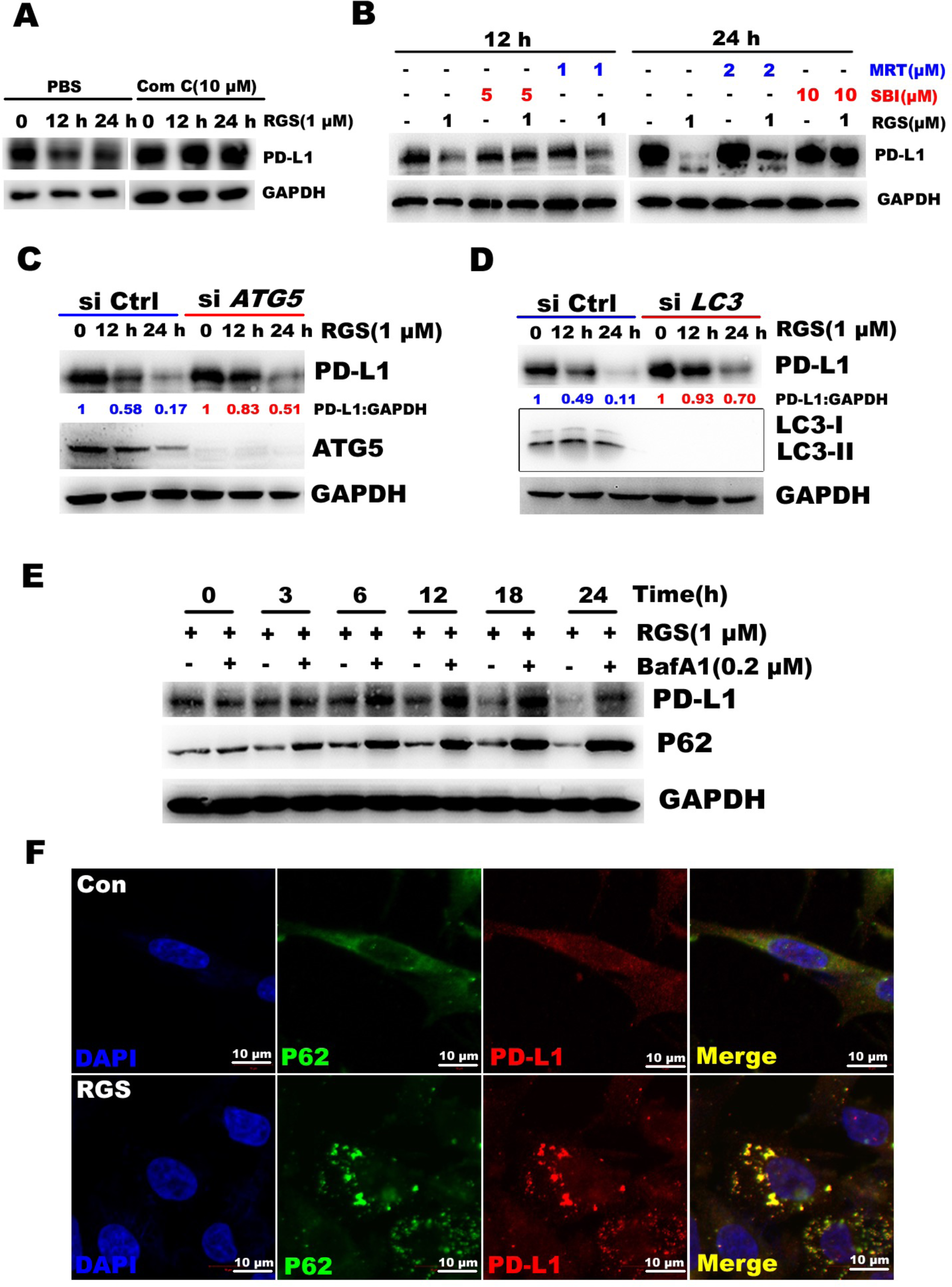
RGS-induced PD-L1 degradation is an autophagy-dependent process. (**A**) RKO cells were pretreated with Compound C for 30 min to inhibit the activity of AMPK. Cells were then treated with or without RGS (1 μ) for 12–24 h, after which the levels of PD-L1 were determined by western blotting. (**B**) RKO cells were pretreated with indicated concentrations of MRT68921, SBI0206965 for 30 min to inhibit the activity of ULK1. Cells were then treated with or without RGS (1μM) for 12–24 h, after which the levels of PD-L1 were determined. (**C and D**) Following transfection with specifically targeted siRNA (*ATG5* or *LC3*) for 24 h, RKO cells were treated with RGS (1 μM) for 0–24 h, after which the levels of PD-L1, ATG5, and LC3 were measured by western bloting; levels of PD-L1 and GAPDH were quantitated using the Image J software. (**E**) RKO cells were pretreated with BafA1 for 30 min to inhibit lysosome function. Cells were then treated with or without RGS (1 μM) for 0–24 h, after which the levels of PD-L1 and P62 were determined by western blotting. (**F**) RKO cells were exposed to RGS (1 μ) for 9 h, and then the distribution of PD-L1 and P62 was analyzed by confocal microscopy. Scale bar, 10 μ

Recent studies have revealed that autophagic receptors, such as SQSTM1/P62 (sequestosome 1), recognize protein aggregates and then bind to LC3 on autophagosomes, thereby delivering the cargos and receptors themselves for degradation [23]. By detecting the expression dynamics of PD-L1 and P62 in RGS-treated RKO cells, we found that, similar to PD-L1, P62 degradation was induced by RGS, whereas, this process was completely blocked by pretreatment with BafA1 (Fig. 6E). Therefore, we hypothesized that P62 served as an autophagic receptor necessary for the degradation of PD-L1. By immunofluorescence, we observed that the co-localization of PD-L1 and P62 was enhanced after treatment with RGS (Fig. 6F). Meanwhile, we observed that knockdown of *P62* evidently impaired the RGS-induced degradation of PD-L1 (Fig. S6C).

Collectively, these results suggested that the RGS-induced degradation of PD-L1 is an autophagy-dependent process, with P62 involved in the process.

### Combination of RGS and cytotoxic T-lymphocyte-associated protein 4 (CTLA-4) blockade effectively suppressed tumor growth in vivo

Recently, combination treatment targeting PD-L1 and CTLA-4 was used in patients with cancer to improve the anti-tumor T cell immunity. Since we showed that RGS could degrade PD-L1, we selected CTLA-4 blockade for combination therapy with RGS in the CT26 CRC mouse model. Immune competent BALB/c mice were treated with RGS and CTLA-4 mAb as indicated in Fig. S7A. As shown in the results, similar to results shown in Fig. 1, RGS alone significantly decreased tumor growth at day 13 compared with the control group (mean tumor weight: 0.388 vs 1.114 g; *P* < 0.01), while a combinatorial treatment of RGS and CTLA-4 mAb exhibited better efficacy (mean tumor weight: 0.101 vs 1.114 g; *P* < 0.01; Fig. 7A–C). Meanwhile, similar to results shown in Fig. S1B, no obvious evidence of toxicity caused by RGS alone or along with CTLA-4 mAb (the combination therapy) was observed, as indicated by the changes in body weight of mice (Fig. S7B). Consistent with our mechanistic findings, results of the immunofluorescence analysis showed that RGS or RGS + CTLA-4 mAb treatment significantly decreased PD-L1 expression, increased infiltration by CD8+CTLs, and enhanced their activity in the tumor xenograft (Fig. 7D–G and S7C–F). These findings suggested that RGS could enhance the efficacy of CTLA-4 mAb therapy in the treatment of CRC.

**Figure 7.**
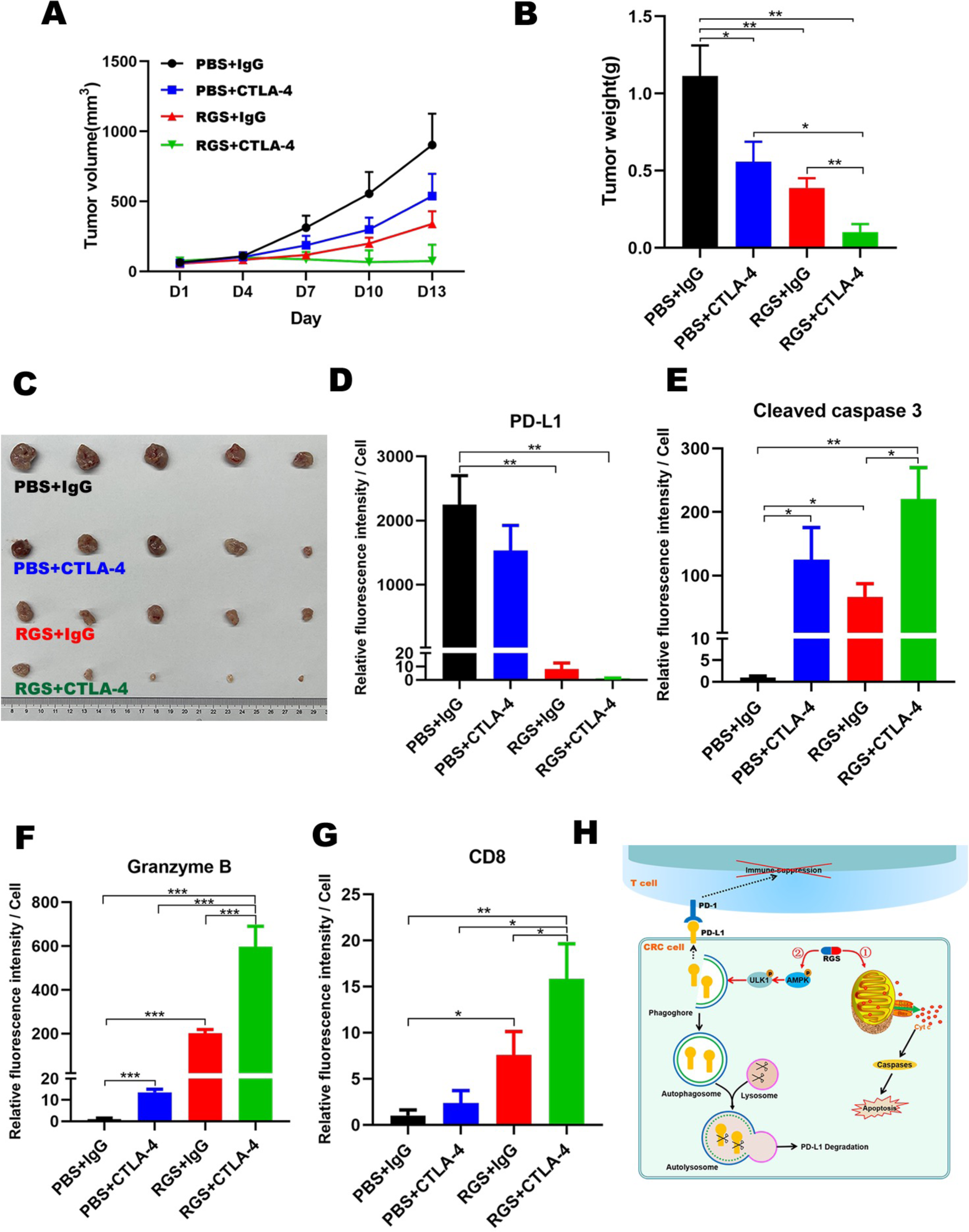
The combination of RGS and CTLA-4 blockade effectively suppressed tumor growth in vivo. (**A**) Anti-tumor effects of RGS, CTLA-4 mAb, and the combinatorial treatment (RGS + CTLA-4 mAb) were assessed in CT26 xenograft models based on BALB/c mice. Tumor volumes were measured every 3 d after palpable tumors reached volumes of 50–100 mm^3^. Tumor growth was evidently inhibited in the combinatorial treated group (N = 5). (**B**) CT26 tumor weight was measured at day 13 (N = 5). (**C**) Representative images of CT26 tumors after RGS and/or CTLA-4 mAb treatment at the end of time point. (**D–G**) Immunostaining, followed by quantification of PD-L1, cleaved caspase 3, granzyme B, and CD8 in the CT26 xenograft tumor based on BALB/c mice using Image J (N = 5). Data shown are mean ± SEM. \**P* < 0.05; \*\**P* < 0.01; \*\*\**P* < 0.001. (**H**) Schematic diagram depicting the mechanism by which RGS exerts its anti-tumor effects. RGS triggers the mitochondria-related apoptosis in CRC cells. Concomitantly, RGS activates AMPK-ULK1 axis dependent autophagy, which plays an essential role in the degradation of PD-L1.

## Discussion

In this study, surprisingly, we elucidated the more evident tumor growth arrest and more robust anti-tumor immunity in the RGS-treated CT26 CRC xenograft tumors based on immune competent mice. Mechanistically, we demonstrated for the first time that RGS inhibited the expression of PD-L1 via autophagy, which was dependent on the activation of AMPK-ULK1 axis. We showed that, preclinically, RGS exhibits synergistic effects with CTLA-4 mAb in the treatment of CRC in immune competent mice.

As a promising anti-cancer drug, RGS has been studied for more than 10 years. Numerous preclinical studies have proved its efficacy in leukemia [10], high-risk myelodysplastic syndrome [24], and solid malignant tumors, such as pancreatic cancer [25], and head and neck squamous cell carcinomas [26]. However, majority of these studies were performed using immune-deficient models, and few studies have focused on the potential role of RGS in modulating tumor immune microenvironment. Recently, in melanoma, Yan et al. reported that RAS-RAF-PI3K pathway inhibition by RGS could induce CD40 expression and activate the immune microenvironment [12]. In the present study, in CRC, besides the elevation of CD40 protein expression, we elucidated that RGS could promote anti-tumor immunity by inducing autophagic degradation of PD-L1.

To investigate the mechanisms underlying the autophagic degradation of PD-L1, we first investigated the phosphorylation of the RAS-RAF-MEK-ERK pathway in RGS-treated RKO cells. Unlike the RAS-mutated DLD1 and HCT116 cells we previously reported, RGS could not block the activation of this pathway in RKO cells.

According to their genetic background, RKO cells harbor BRAF ^V600E^ mutation, which could constitutively activate the signaling pathway without upstream stimulation [27]. Therefore, in RKO cells, the RAF downstream MEK-ERK activation could not be prohibited even when RAS-RAF interaction were blocked by RGS [28]. Considering the role of RAF-MEK-ERK activation in stabilizing PD-L1 mRNA [5], we correspondingly confirmed that the expression of PD-L1 mRNA was not altered in RKO cells after 0–12 h of incubation with RGS (Fig. 3A). Thus, the inhibitory effect of RGS on the expression of PD-L1 was probably independent of blocking the RAF-MEK-ERK activation or reducing the level of PD-L1 mRNA.

Autophagy involves the turnover of protein aggregates, cytoplasmic macromolecules, and defective organelles by sequestration via double-layered membranes that form autophagosomes, which eventually deliver the contents to lysosomes where “cargos” are degraded. Wani et al. reported that autophagy induced by alborixin, an ionophore, substantially cleared Aβ in microglia and primary neurons β [29]. Liu et al. demonstrated that a novel hypoxia-activated prodrug, Q6, degraded hypoxia inducible factor 1A via an autophagy-dependent mechanism in hepatocellular carcinoma [30]. Therefore, autophagy induced by small-molecule drugs might down-regulate crucial proteins, such as PD-L1, in cancer therapies. In our study, we confirmed that RGS induced the autophagy-dependent degradation of PD-L1. Furthermore, by comparing the expression of PD-L1 and other common membrane proteins in CRC cells under incubation with RGS, we found that, unlike HGFR and E-cadherin, autophagic degradation targeting PD-L1 was a specific process. This specificity could be attributed to the participation of P62 in the autophagic degradation process. P62 has been elucidated to be a selective autophagic receptor, which can link polyubiquitinated proteins to autophagosome membranes and deliver the aggregates for degradation [23]. In our study, we found that RGS induced a significant co-localization between PD-L1 and P62, whereas knocking down the expression of *P62* impaired the RGS-induced degradation of PD-L1.

In addition to the regulatory effects of RGS on PD-L1, the cytotoxic effects of RGS in RKO cells in vivo and in vitro were investigated. Similar to our previous findings [11], RGS induced mitochondria-related apoptosis in a time- and dose-dependent manner (Fig. S3).

In conclusion, we propose a more nuanced conceptual model explaining the anti-tumor activity of RGS (Fig. 7H). In this model, RGS triggers two mutually independent events: initially, RGS induces cancer cell mitochondria-related apoptosis, and then provokes the autophagy-dependent degradation of tumor PD-L1, which plays a pivotal role in strengthening anti-tumor immunity.

Our study had few limitations. First, in contrast to the multiple CRC cell lines (DLD1, LOVO, RKO, HCT116, and CT26) used to confirm the PD-L1 inhibitory effect of RGS, the majority of experiments dissecting the underlying mechanism were performed in RKO cells, the only CRC cell line that expressed endogenous PD-L1. Second, RGS exhibited impressive cytotoxic effects in RKO cells, which harbored BRAF ^V600E^ mutation. Considering the minimal inhibition in RAS-BRAF ^V600E^ -MEK-ERK activation, the cytotoxic effects seemed to be independent of the inhibition of this pathway. This interesting phenomenon might be, at least partly, explained by the potential RGS-induced decline in CRAF phosphorylation [28]. Mielgo et al. reported that, inhibition of CRAF^S338^ phosphorylation, not BRAF, inhibited the association of CRAF with polo-like kinase 1, leading to mitotic progression block and subsequent apoptosis [31]. Thus, as shown in Fig. S4A, without evident decrease in the levels of p-MEK or p-ERK, mitosis-specific histone H3 (Ser10) phosphorylation was elevated after 6 h of treatment with RGS ahead of apoptosis protein markers. Nevertheless, the other reported cytotoxic mechanisms of RGS, including the inhibition of polo-like kinase 1 activity [32] or microtubule destabilization [33], were not investigated in our study; our results could not exclude the potential contribution of these mechanisms in the observed cytotoxicity.

In conclusion, our findings showed that, apart from the impressive cytotoxicity, RGS induced autophagy-dependent degradation of tumor PD-L1, which strengthened anti-tumor immunity in CRC. We proved that RGS could improve response to immune checkpoint blockade in CRC. Therefore, our findings provide a theoretical basis for conducting further clinical trials to investigate the efficacy of RGS and its combination with immune checkpoint inhibitors.

## Materials and Methods

### Chemicals and reagents

RGS (S1362), CQ (S6999), BafA1 (S1413), MG132 (S2619), CHX (S7418), MRT68921 (S7949), Compound C (S7840), and SBI0206965 (S7885) were purchased from Selleck Chemicals Company (Houston, TX, USA). Rapamycin (S1842) and Z-VAD-FMK (C1202) were purchased from Beyotime Biotechnology (Shanghai, China). LysoTracker RED (40739ES50) was obtained from Yeasen Biotechnology (Shanghai, China). All chemicals were dissolved in Dimethyl Sulfoxide (DMSO) (Sigma-Aldrich, D8418) and stored at –20

### Cell culture

Human CRC cell lines (HT29, DLD1, HCT116, SW480, SW620, RKO, LOVO, SW48, and Caco-2) and murine CT26 CRC cells were obtained from the American Type Culture Collection (Rockville, MD, USA) in October, 2016. Following receipt, cells were grown and frozen as seed stocks. Cells were passaged for a maximum of 3 months, after which new seed stocks were thawed. Cell lines were authenticated using DNA fingerprinting (variable number of tandem repeats). Caco-2 cells were cultured in minimal essential medium (Gibco, 10370070) supplemented with 20% fetal bovine serum (FBS) (Gibco, 10099), while other cells were maintained in RPMI 1640 medium (Gibco, 11875119) supplemented with 10% FBS, 100 units/mL penicillin, and 100 mg/mL streptomycin at 37[in a humidified atmosphere of 5% carbon dioxide. All cell lines were routinely screened for the presence of mycoplasma (Mycoplasma detection kit, Sigma-Aldrich, MP0035).

### Western blot assay

Total protein was extracted from CRC cells after treatment with RGS, and protein concentration was determined using the Pierce™ BCA Protein Assay kit (Thermo Fisher Scientific, 23225). Protein samples were subjected to 10–12% SDS-PAGE and transferred to a polyvinylidene fluoride membrane (Bio-Rad, Hercules, CA, USA).

After blocking the membranes with 5% nonfat milk for 1 h at 25 [we incubated the membranes with appropriate primary antibodies overnight at 4 [Primary antibodies against human PD-L1 (13684), HGFR (8198), E-cadherin (14472), cleaved caspase 3 (9664), cleaved caspase 9 (20750), PARP (9532), LC3 (3868), p-MEK1 (S217/221) (9154), p-ERK1/2 (T202/204) (4370), p-AKT (S473) (4060), p-mTOR (S2448) (5536), p-AKT (T308) (13038), AMPK (5832), p-AMPK (T172) (50081), p-ULK1 (S757) (14202), ERK1/2 (4695), ULK1 (8054), and AKT (4691) were purchased from Cell Signaling Technology. All mentioned antibodies were used in a dilution of 1:1000. Primary antibodies against ATG5 (ab108327), murine PD-L1 (ab213480), p-ULK1 (S555) (ab133747), and P62 (ab109012) (all diluted 1:1000) were purchased from Abcam. Primary antibodies against MEK1/2 (AF1057), and p-histone H3 (S10) (AF 1180) (all diluted 1:1000) were purchased from Beyotime Biotechnology. Α mouse anti-glyceraldehyde 3-phosphate dehydrogenase (GAPDH) monoclonal antibody (dilution 1:1000; Cell Signaling Technology, 5174) was used as the loading control. Following incubation with a secondary antibody conjugated to horseradish peroxidase for 1 h at 25[, immunoreactive bands were visualized using an enhanced chemiluminescence detection system (Thermo Fisher Scientific, 32109).

### Mouse xenograft CRC model

All animal procedures were performed in accordance with protocols reviewed and approved by the Animal Ethics Committee of the Second Affiliated Hospital of Zhejiang University School of Medicine (approval no. 2020.035). Briefly, 5- to 6-week-old female nude mice and BALB/c mice were purchased from SLAC Laboratory Animal Co., Ltd. (Shanghai, China). RKO (1 × 10^6^) and CT26 (5 × 10^5^) cells were suspended in PBS or Matrigel (Corning, 354230) and then subcutaneously injected into mice. Tumor growth was monitored daily until the tumor was palpable (50–100 mm^3^). Subsequently, mice were randomized into groups, with each group receiving either PBS or RGS (100 mg/kg/day) or CTLA-4 mAb (10 mg/kg, twice a week; Bio X Cell, BE0032) or corresponding isotype (Bio X Cell, BE0091) control via intraperitoneal injection. Body weight and tumor size were measured every 3 d. Once the tumor size reached 15–20 mm in any dimension or animals became ill, tumor fragments were harvested. Tumor volumes were calculated in mm^3^ using the following formula:

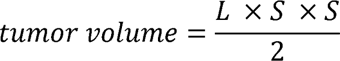

where *L* is the long axis of the tumor, and *S* is the short axis of the tumor. After 2 weeks of administration, mice were sacrificed, and tumors were excised, photographed, fixed in 10% neutral formalin, and embedded in paraffin.

### CyTOF analysis of immune cells

CyTOF analysis was performed by PLT-Tech Inc. (Hangzhou, China) according to a previously described protocol [34]. In brief, tumor tissue was dissociated into single cells with DNAse (Sigma-Aldrich, AMPD1), collagenase IV (Sigma-Aldrich, C4-BIOC), and hyaluronidase (Sigma-Aldrich, H1115000). The digested cells were resuspended and stained for viability with Pt-194 (Fluidigm, 201194). Qualified samples were blocked with Fc receptors and stained for 30 min with surface metal-labeled monoclonal antibodies. Then, the cells were incubated overnight with DNA Intercalator-Ir (Fluidigm, 201192A) to discriminate singly nucleated cells from doublets. Permeabilization buffer was applied, and the cells were incubated in an intracellular antibody panel. Next, the cells were rinsed, and the signals were detected using a CyTOF system (Helios™, Fluidigm). Immune cell types were identified via nonlinear dimensionality reduction (t-distributed stochastic neighbour embedding), followed by density clustering.

### Transmission electron microscopy

Cells were fixed in 2.5% glutaraldehyde solution (Sigma-Aldrich, G6257) for 1 h at 25°C. Then, cells were harvested, washed gently three times in PBS, and fixed in 1% osmium tetroxide (Sigma-Aldrich, 75632) for another 1 h. After dehydration in a graded ethanol series, cells were critical-point dried and sputter-coated with 10% gold. Cells were observed under a transmission electron microscope (TECNAI10, Philips, Netherlands).

### siRNA transfection

Cells were seeded on 6-well plates and transfected 24 h later using Lipofectamine 3000 (Invitrogen, L3000150), according to the manufacturer’s instructions. Human *LC3* siRNA, *ULK1* siRNA, *P62* siRNA, *ATG5* siRNA, and control siRNA were purchased from Tsingke Biological Technology (Hangzhou, Zhejiang, China). The siRNA sequences were as follows: *LC3*, 5′- GGUGAGAAGCAGCUUCCUGUUCUGGAUAA -3′; *ULK1-1*, 5′- CUUCCAGGAAAUGGCUAAUUCUGUCUACC -3′; *ULK1-2*, 5′- GUACCUCCAGAGCAACAUGAUGGCGGCCA -3′; *P62-1*, 5′-AGACUACGACUUGUGUAGCGUCUGCGAGG -3′; *P62-2*, 5′- CGGCAGAAUCAGCUUCUGGUCCATCGGAG -3′; *ATG5,* 5′- GUGCUUCGAGAUGUGUGGUUUGGACGAAU -3′.

### Immunofluorescence

RKO cells were planted on coverslips and treated with RGS for the indicated times. Cells were fixed with 4% (w/v) paraformaldehyde and permeabilized with PBS containing 0.5% Triton X-100. After blocking the cells with 5% bovine serum albumin, we stained the cells with the following primary antibodies: anti-LC3 antibody (1:500; Cell signaling technology, 3868), anti-PD-L1 antibody (1:500; Proteintech, 66248-1-Ig), and anti-P62 antibody (1:500; Abcam, ab109012) at 4 [overnight. After washing the cells twice with PBS containing 0.05% Tween 20, we incubated the cells with the appropriate Alexa Fluor 488 and 555 secondary antibodies (Invitrogen, A-11008, A32727) in the dark. Nuclei were visualized by staining with 46-diamidino-2-phenylindole (Sigma-Aldrich, P36980). Images were obtained using a confocal microscope (Zeiss, LSM 710). Image analysis was performed using the Zen software.

### Statistical analysis

All data are presented as the mean ± standard error of the mean of 3 independent experiments. Statistical analysis was performed using Student’s *t*-test or analysis of variance with multiple comparisons using the Prism software (version 6.0; GraphPad, La Jolla, CA, USA). Differences were considered significant at *P* < 0.05, *P* < 0.01, and *P* < 0.001. A full description of additional materials and methods employed in our study is provided as Supplemental Information.

### Data availability statement

The data sets used for the current study are available from the corresponding author (the Lead Contact, Qian Xiao) upon reasonable request.

## Supporting information

Supplemental Experimental Procedures and Supplemental Figures and Legends

## Acknowledgments

We would like to thank Dr. Jinlong Tang and Dr. Qi Yang (from the Department of Pathology, the Second Affiliated Hospital of Zhejiang University School of Medicine) for their generous technical assistance in immunohistochemical staining.

