## Supplemental Experimental Procedures and Supplemental Figures and Legends for "Rigosertib promotes anti-tumor immunity via autophagic degradation of PD-L1 in colorectal cancer cells"

**Supplemental Information**

**Supplemental Experimental Procedures**

**Cell viability analysis**

Cell viability was analyzed using the Cell Counting Kit 8 (CCK8; Dojindo Laboratories, CK04). Cells were seeded into 96-well plates at a density of 5–10 × 10^3^ cells per well overnight. The RGS working solution was diluted with complete medium with a maximal concentration of 0.1% DMSO. Cells were treated with the indicated concentrations of RGS. After incubation, CCK8 was added to each well. Absorbance was measured using a microplate reader at 450 nm after incubation for an additional 2 h. Three replicate wells were set up for each group.

**Plasmid construction**

The pLV-EGFP-N lentiviral vector (Yingmaoshengye Biotechnology, Beijing, China) and XhoI and EcoRI restriction endonucleases were used for plasmid construction. Vector and target genes were digested at 37°C for 0.5 h, then subjected to agarose gel electrophoresis, and recovered. Vector and target genes were mixed at a ratio of 1:7 and ligated with T4 ligase at 25°C overnight. Next, acquired lentiviral constructs were used to transform DH5α competent cells. After mixing, the bacterial solution was collected on an LB plate containing ampicillin. Plates were placed in a 37°C-incubator for 12–16 h, and a single colony was picked from each plate for reproduction, sequencing, and identification.

**Lentiviral transduction and generation of stable cell lines**

Lentiviruses were produced by transfecting HEK293T cells with the psPAX2 packaging plasmid (Addgene plasmid, 12260), pMD2.G envelope plasmid (Addgene plasmid, 12259), and PD-L1-EGFP plasmids, using pLVX-IRES-puro (Youbio, VT1464) as the control. Cell supernatants were collected at 24 and 48 h after transfection and were used for infection or stored at –80°C. To obtain stable cell lines, we infected cells at 70–80% confluence with lentivirus diluted 1:1 with normal cell culture medium in 96-well plates for 24 h. After 24 h of infection, the supernatants were replaced with normal cell culture medium. After 48 h, cells were subjected to puromycin selection for approximately 1 week in 24-well plates and passaged before use. Puromycin was used at a concentration of 2 µg/mL to maintain RKO cells.

**Mitochondrial fractionation and analysis**

Mitochondria-enriched fractionation was performed using a Mitochondria isolation kit (Thermo Fisher Scientific, Waltham, MA, USA) according to the manufacturer’s instructions. The fractions of RGS treated RKO cells were examined by western blot assay using anti- Bcl-2-associated X protein (Abcam, ab32503), anti- cytochrome c (Abcam, ab133504), anti- cytochrome c oxidase IV (Abcam, ab202554), and anti-GAPDH (Cell Signaling Technology, 5174) antibodies.

**Immunohistochemistry staining**

Paraffin-embedded tissue sections were immunostained using a two-step method. Following deparaffinization with xylene, rehydration through graded ethanol, and antigen retrieval using EDTA (100°C for 15 min), the endogenous peroxidase activity and nonspecific antigen-binding sites were blocked by successive incubation with 3% hydrogen peroxide for 10 min and 5% bovine serum albumin (Bio-Rad Laboratories, 5000206) for 30 min at 25°C. Tissue sections were then incubated with rabbit anti-PD-L1 monoclonal antibody (1:200; Cell Signaling Technology, 13684S) at 37°C for 2 h. Bound antibodies were detected using the REAL™ EnVision™ detection system (Agilent Technologies, K5007), and sections were counterstained with hematoxylin for 2 min at 25°C. Negative controls were prepared using the same procedure without primary antibody staining. Immunohistochemical staining was reviewed and scored by 2 independent pathologists blinded to the study.

**qRT-PCR assay**

Total RNA from cells was isolated using TRIzol reagent (Invitrogen, 15596026) according to the manufacturer’s instructions. The quality and quantity of RNA were evaluated using a NanoDrop 2000 spectrophotometer (Thermo Fisher Scientific, Pittsburgh, PA, USA). cDNA was synthesized using the PrimeScript™ II 1st strand cDNA synthesis kit (Takara Biotechnology, 6210A). Quantitative real-time RT-PCR was performed using a standard SYBR-Green PCR kit protocol (YEASEN, Shanghai, China) with a 7500 Fast Real-Time PCR system (Life Technologies, Shanghai, China). Gene expression was quantified by the Taqman probe system using the following primers: *PD-L1* forward: 5′-TGCCGACTACAAGCGAATTACTG-3′, reverse: 5′-CTGCTTGTCCAGATGACTTCGG-3′; *GAPDH*, forward: 5′-AGAAGGCTGGGGCTCATTTG-3′, reverse: 5′-AGGGGCCATCCACAGTCTTC-3′. All PCR reactions were performed in triplicate. Gene expression was normalized to the expression of *Gapdh*.

**mRFP-GFP-LC3 transfection**

mRFP-GFP-LC3 vectors were transfected according to the manufacturer’s instructions (Youbio Biotechnology, China), and transfected cells were cultured for 72 h at 37°C and 5% carbon dioxide.

**Immunofluorescence staining of frozen sections**

Tumor masses were resected from mice and fixed with 4% paraformaldehyde overnight. Tumor masses were dehydrated in 30% sucrose, embedded in OCT block, and frozen for cryostat sectioning. After washing the sections with PBS, we incubated cryostat sections (8-mm-thick) in blocking solution (5% BSA containing 0.2% Triton X-100, pH 7.4) for 40 min at 25°C. Samples were stained with primary antibodies against cleaved caspase 3 (1:400; Cell Signaling Technology, 9661), CD8 (1:100; BioRad, MCA609G,), Granzyme B (1:200; Cell Signaling Technology, 17215), PD-L1 (1:200; Abcam, ab213480) overnight at 4°C, followed by incubation with AlexaFluor 488 and 555 (Abcam, ab150077, ab150158, ab150078) secondary antibodies at 25°C for 1 h. DAPI Fluoromount-G (SouthernBiotech, 0100-20) was used for nuclear staining.

**Supplemental Figures**

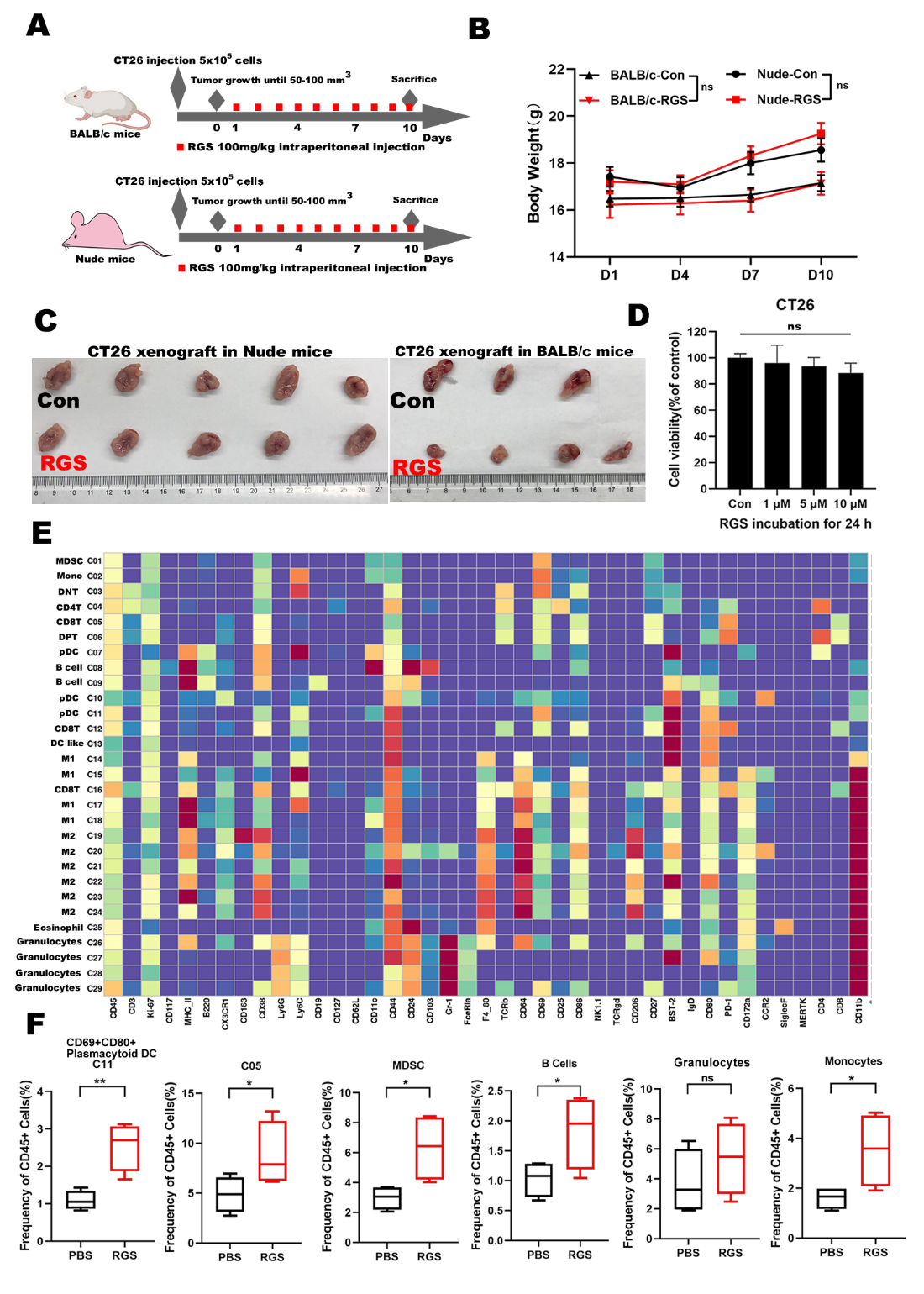

**Figure S1.** RGS reprogramed the tumor immune environment in colorectal cancer. (**A and B**) In the CT26 xenograft models based on nude and BALB/c mice, no significant difference was observed in the weight of animals between PBS- and RGS-treated groups. (**C**) Representative images of tumor gross morphology in CT26 xenograft models based on nude or BALB/c mice. (**D**) The cell viability of murine CT26 cells treated with different concentrations of RGS for 24 h was assessed by CCK8 assay.

(**E**) Heatmap showed the differential expressions of 41 immune markers in the 29 cell clusters. The label on the left shows the immune cell types of clusters according to typically expressed markers. (**F**) Data derived from CyTOF. Frequencies of C11, C05, myeloid-derived suppressor cells (MDSCs), B cells, granulocytes, and monocytes between PBS and RGS treated tumors. Data shown are mean ± SEM. **P* < 0.05.

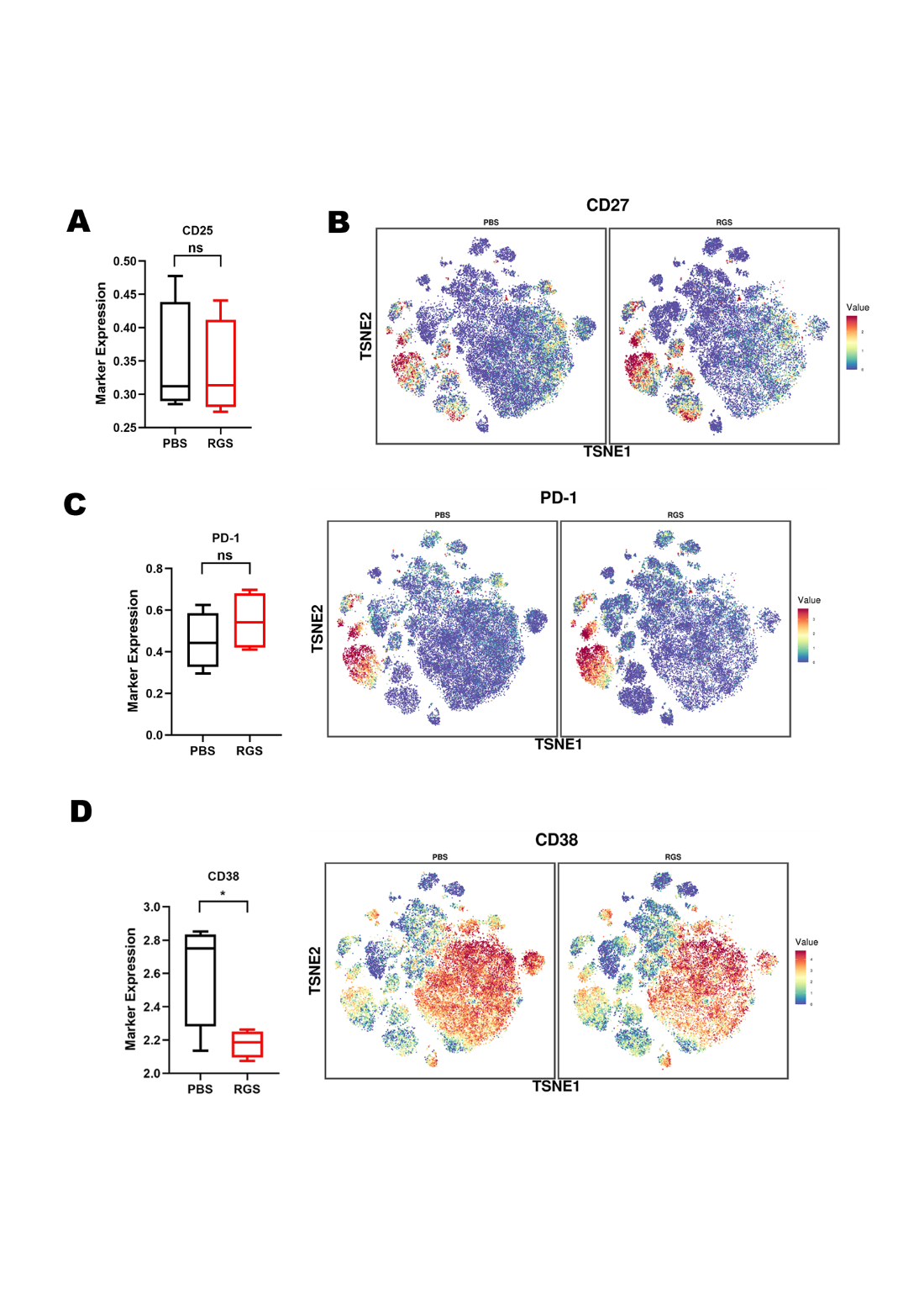

**Figure S2.** Difference in expression of functional molecules between PBS- and RGS-treated CT26 xenograft tumors based on BALB/c mice. Box plot and t-SNE plot showed the differential expressions of CD25 (**A**), CD27 (**B**), PD-1 (**C**) and CD38 (**D**) between PBS- and RGS-treated CT26 tumors. Data derived from CyTOF. Data shown are mean ± SEM. **P* < 0.05.

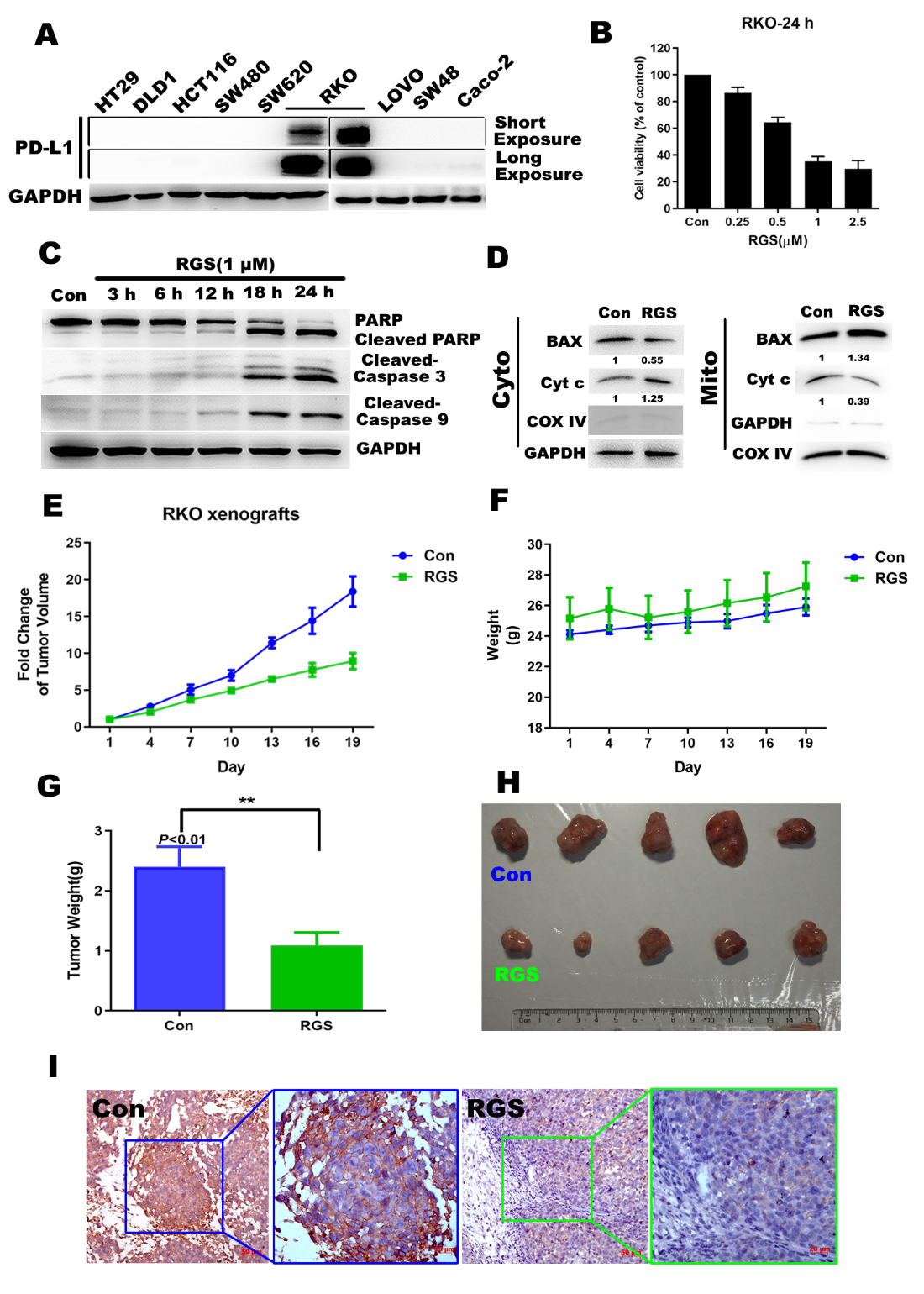

**Figure S3.** RGS showed direct cytotoxicity towards RKO cells both in vitro and in vivo. (**A**) Level of PD-L1 protein in HT29, DLD1, HCT116, SW480, SW620, RKO, LOVO, SW48, and Caco-2 CRC cells was evaluated by western blotting. GAPDH was used as loading control. (**B**) The viability of RKO cells following treatment with different concentrations of RGS for 24 h was assessed using the CCK8 assay. (**C**) RKO cells were treated with RGS for various time periods prior to lysis, and apoptosis-related protein markers were examined as indicated. (**D**) Levels of apoptosis-related proteins (Bcl-2-associated X protein [BAX] and cytochrome c [Cyt c]) were determined by western blot assay of cytoplasmic and mitochondrial fractions of RKO cells following 24 h incubation with RGS. Cytochrome c oxidase IV (COX IV) was used as the loading control for the mitochondrial fraction. (**E**) The cytotoxicity of RGS was assessed in RKO xenograft mouse models. Tumor volumes were measured every 3 d after palpable tumors reached 50–100 mm^3^. Tumor growth was significantly inhibited in the RGS-treated RKO xenograft model (N = 5). (**F**) No difference was observed in the weight of animals between PBS- and RGS-treated nude mice. (**G**) Tumor weight was assessed on sacrificed mice, suggesting a significant decrease in RGS-treated mice in the RKO xenograft model. (**H**) Representative images of gross morphology. (**I**) The level of PD-L1 in the subcutaneous RKO xenograft tumors were evaluated by immunohistochemical staining. Data are presented as the mean ± SEM obtained from at least three independent experiments performed in triplicate. **P* < 0.05; ***P* < 0.01.

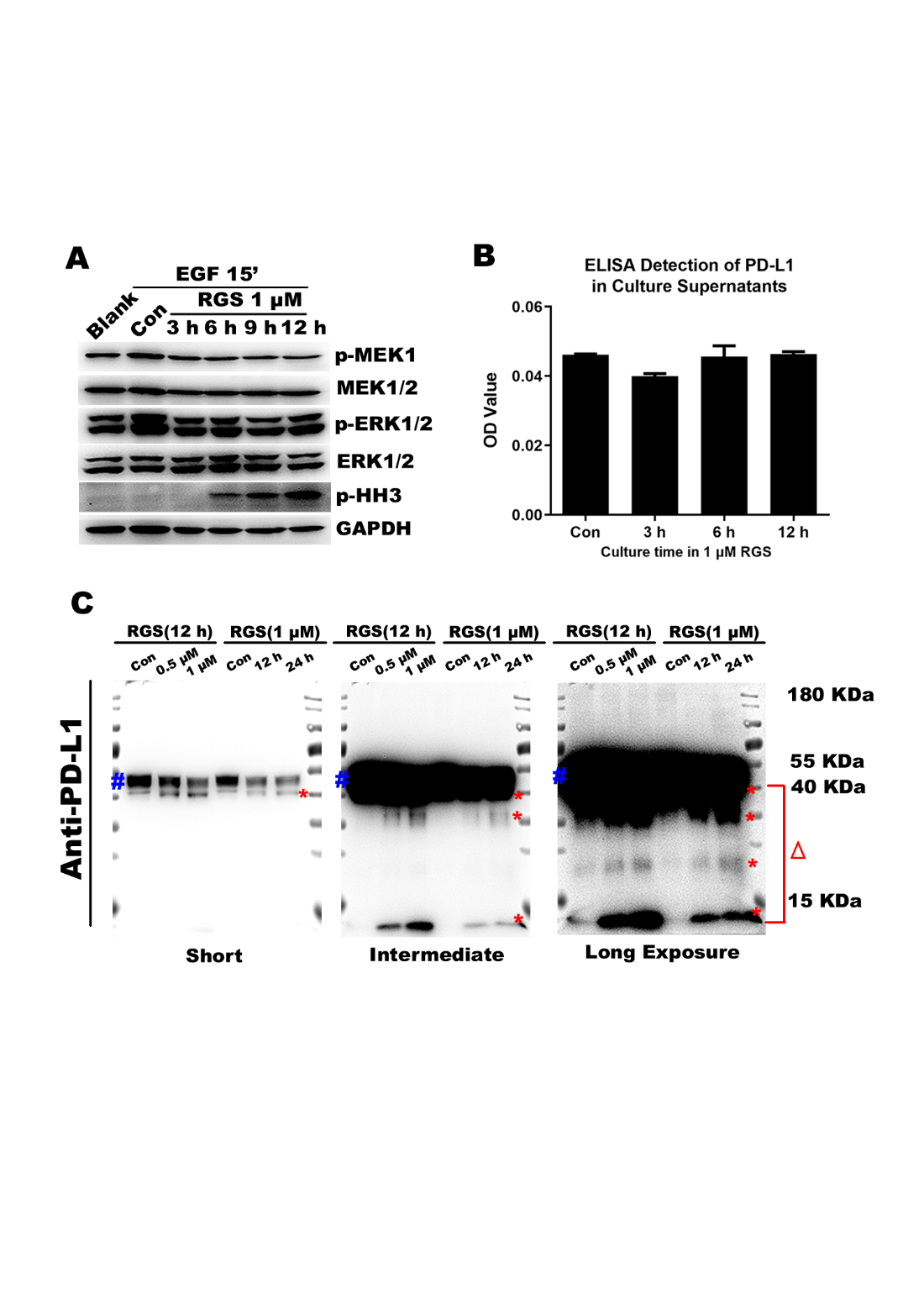

**Figure S4.** RGS decreased the level of PD-L1 by accelerating its degradation. (**A**) RKO cells were serum starved overnight and treated as indicated with DMSO or 1 μM RGS prior to stimulation with EGF for 15 min and subsequent lysis. Lysates were examined for levels of MEK, p-MEK, ERK, p-ERK, and p-histone H3 (HH3) by western blotting. (**B**) RKO cells were treated with 1 μM RGS for 0–12 h. Subsequently, secreted PD-L1 in culture medium supernatants were detected by ELISA. (**C**) Full-length PD-L1 and its degraded fragments were detected using anti-PD-L1 mAb via western blotting in RKO cells after treatment with 0.5–1 μM RGS for 12–24 h. Full-length PD-L1 is indicated by BLUE #, whereas PD-L1 fragments are indicated by RED * and △. Data are presented as the mean ± SEM from three different experiments performed in triplicate.

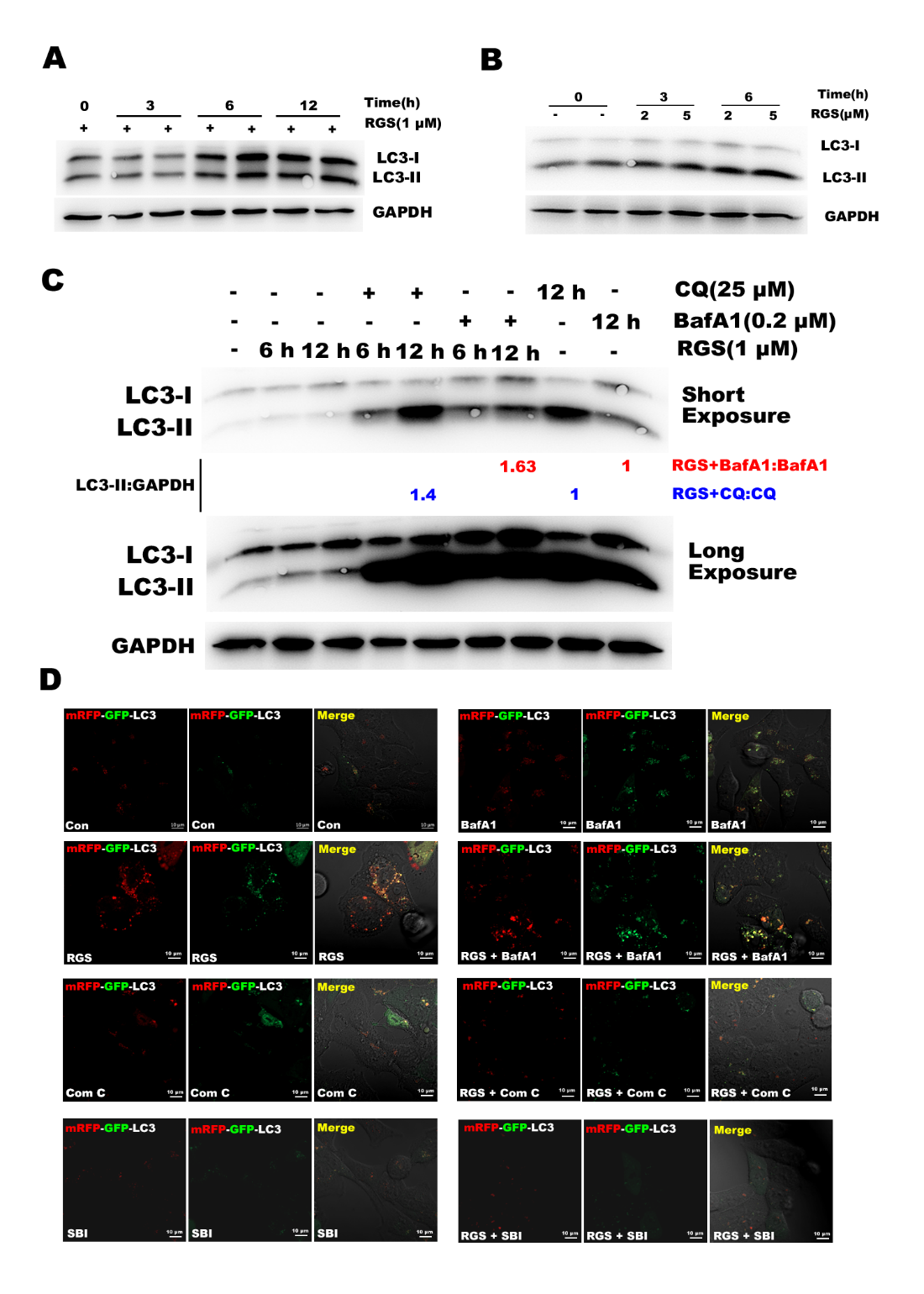

**Figure S5.** RGS induced LC3-II turnover in RKO cells. (**A and B**) RKO cells were treated with 1 μM or 2–5 μM RGS for different time periods, and the levels of LC3 were then measured by western blotting. (**C**) RKO cells were pretreated with CQ or BafA1 for 30 min to inhibit lysosomal function. Cells were then treated with or without RGS (1 μM) for 6–12 h, after which the levels of LC3 were determined by western blotting. The levels of LC3-II and GAPDH were quantitated using the Image J software. (**D**) Representative images of tandem mRFP-GFP-LC3 stably expressing HCT116 cells. HCT116 cells stably transfected with mRFP-GFP-LC3 were pretreated with BafA1, Compound C, or SBI0206965 for 30 min, then the cells were treated with 1 μM RGS for 12 h. Scale bar, 10 μm.

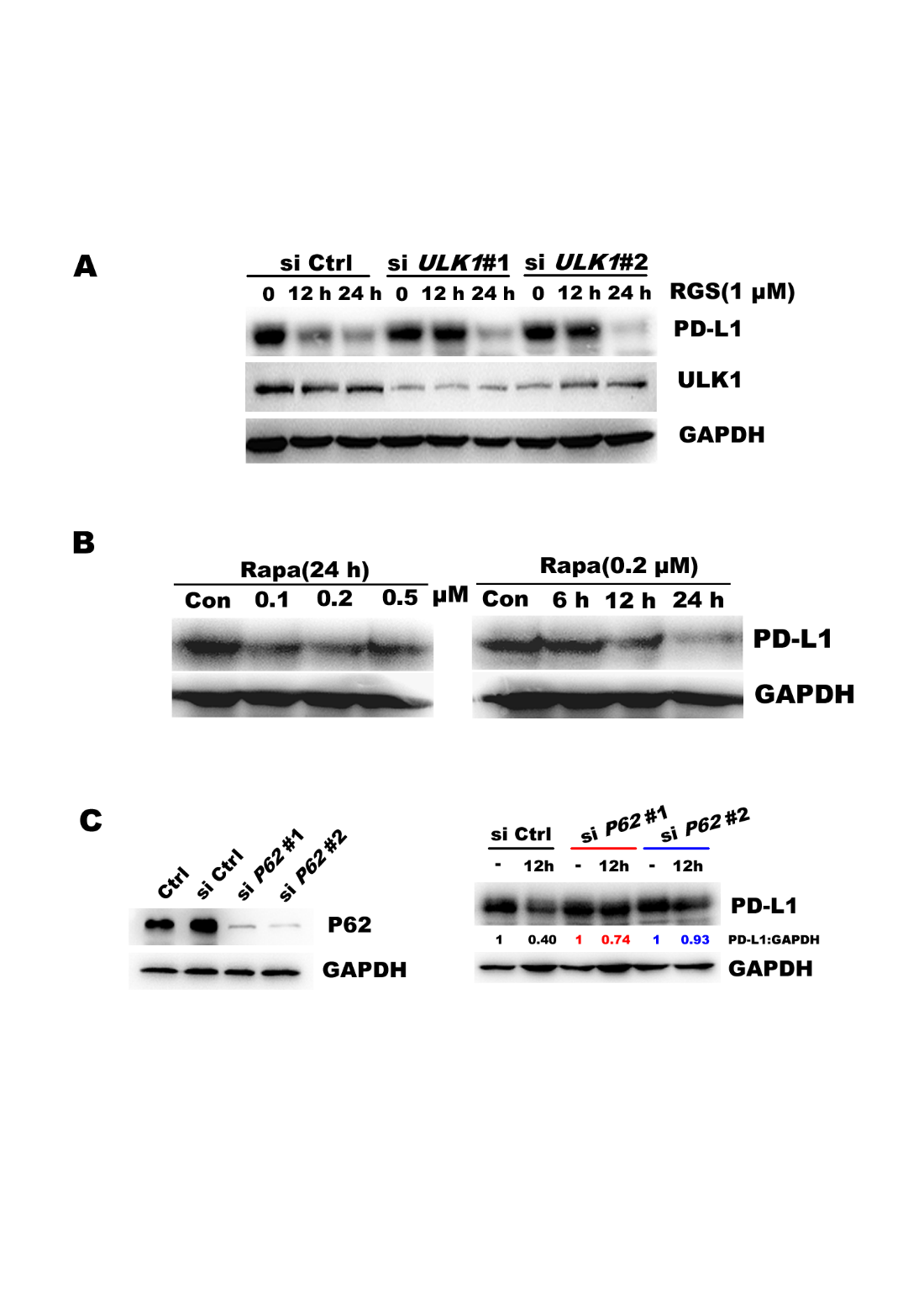

**Figure S6.** Autophagy regulated RGS-induced PD-L1 degradation. (**A**) After transfection with specifically targeted siRNA (si*ULK1*) for 24 h, RKO cells were treated with RGS (1 μM) for 0–24 h. Subsequently, the levels of PD-L1 and ULK1 were measured by western blotting. (**B**) RKO cells were treated with 0.1, 0.2, and 0.5 μM rapamycin for 24 h or 0.2 μM for various time periods prior to lysis. The levels of PD-L1 were examined by western blotting. (**C**) After transfection with specifically siRNA targeting *P62* for 24 h, the remaining levels of P62 in RKO cells were determined. Following knockdown of *P62*, RKO cells were treated with RGS (1 μM) for 12 h, after which the levels of PD-L1 were measured; levels of PD-L1 and GAPDH were quantitated using the Image J software.

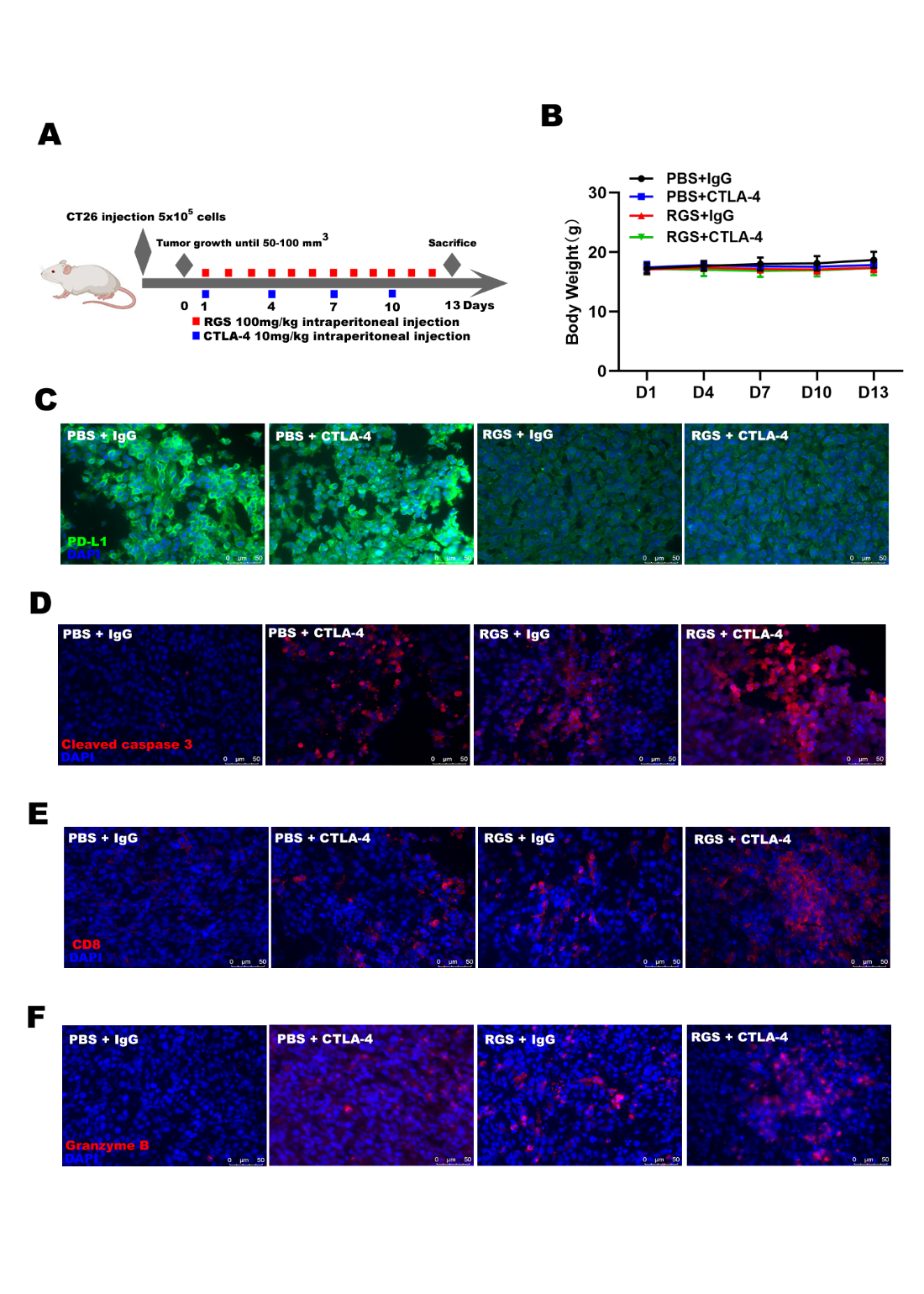

**Figure S7**. The combination of RGS and CTLA-4, effectively suppressed tumor growth in vivo. (**A and B**) In the CT26 xenograft models based on BALB/c mice, no significant difference was observed in the weight of animals between RGS, CTLA-4 mAb, and the combinatorial treatment groups. (**C–F**) Immunostaining of PD-L1, cleaved caspase 3, CD8, and granzyme B in the CT26 xenograft tumors based on BALB/c mice. Scale bar, 50 μm.
